# Pan-cancer scale screening reveals NRF2 associated immunoevasive characteristics in non-small cell lung cancer and squamous malignancies

**DOI:** 10.1101/2022.05.15.489654

**Authors:** Jouni Härkönen, Petri Pölönen, Ashik Jawahar Deen, Ilakya Selvarajan, Hanna-Riikka Teppo, Elitsa Y Dimova, Thomas Kietzmann, Maarit Ahtiainen, Tiia Eklund, Teijo Kuopio, Eva-Maria Talvitie, Pekka Taimen, Markku Kallajoki, Minna U Kaikkonen, Merja Heinäniemi, Anna-Liisa Levonen

**Affiliations:** Faculty of Health Sciences, A.I. Virtanen Institute for Molecular Sciences, University of Eastern Finland, Kuopio, 70210, Finland; Faculty of Health Sciences, Institute of Biomedicine, University of Eastern Finland, Kuopio, 70210, Finland; Cancer Research and Translational Medicine Research Unit, University of Oulu, Oulu, 90220, Finland; Medical Research Center Oulu, Oulu University Hospital and University of Oulu, Oulu, 90570, Finland; Department of Pathology, Oulu University Hospital, Oulu, 90220, Finland; Faculty of Biochemistry and Molecular Medicine, University of Oulu and Biocenter Oulu, Oulu, 90570, Finland; Department of Education and Research, Hospital Nova of Central Finland, Jyväskylä, 40620, Finland; Department of Biological and Environmental Science, University of Jyväskylä, Jyväskylä, 40100, Finland; Department of Pathology, Hospital Nova of Central Finland, Jyväskylä, 40620, Finland; Department of Genomics, Turku University Hospital and University of Turku, Turku, 20520, Finland; Institute of Biomedicine and FICAN West Cancer Centre, University of Turku, Turku, 20520, Finland; Department of Pathology, Turku University Hospital, Turku, 20521, Finland

**Keywords:** NRF2, KEAP1, redox, squamous, NSCLC, T-cells, Interferon gamma, HLA-I, SOX2, TP63

## Abstract

The NRF2 pathway is frequently activated in various cancer types, yet a comprehensive analysis of its effects across different malignancies is currently lacking. We developed a robust NRF2 activity metric and utilized it to conduct a pan-cancer wide analysis of oncogenic NRF2 signaling. We identified a distinct immunoevasive phenotype where high NRF2 activity is associated with low interferon-gamma (IFNγ), HLA-I expression and T-cell infiltration spanning non-small cell lung cancer (NSCLC) and squamous malignancies of head and neck area, cervix and esophagus. In squamous cell cancers, NRF2 overactive tumors comprise a molecular phenotype with *SOX2*/*TP63* amplification, *TP53* mutation and *CDKN2A* loss. These immune-cold NRF2 hyperactive diseases are associated with upregulation of immunomodulatory *NAMPT, WNT5A, SPP1, SLC7A11* and *SLC2A1* that represent candidate NRF2 target genes, suggesting direct modulation of the tumor immune milieu. Based on single-cell mRNA data, coupled with a priori information on intercellular ligand-receptor interactions, cancer cells of this subtype exhibit decreased expression of IFNγ responsive ligands, and increased expression of immunosuppressive ligands *NAMPT, SPP1* and *WNT5A* that mediate signaling in intercellular crosstalk. As we observed differential cytokine mRNA expression with IFNγ treatment in NSCLC adenocarcinoma subtype, we explored the cytokine secretome *in vitro*. We found that secreted neutrophil chemoattractants interleukin-8 (*CXCL8*) and ENA-78 (*CXCL5*) are elevated in NRF2 overactive cells, suggesting contribution of immunosuppressive neutrophils in NRF2 driven immune escape. Importantly, as overactive NRF2 is associated with immune-cold characteristics, our results highlight the utility of NRF2 pathway activation as a putative biomarker for stratifying immune-checkpoint blockade responders and non-responders across NSCLC and squamous cancers.

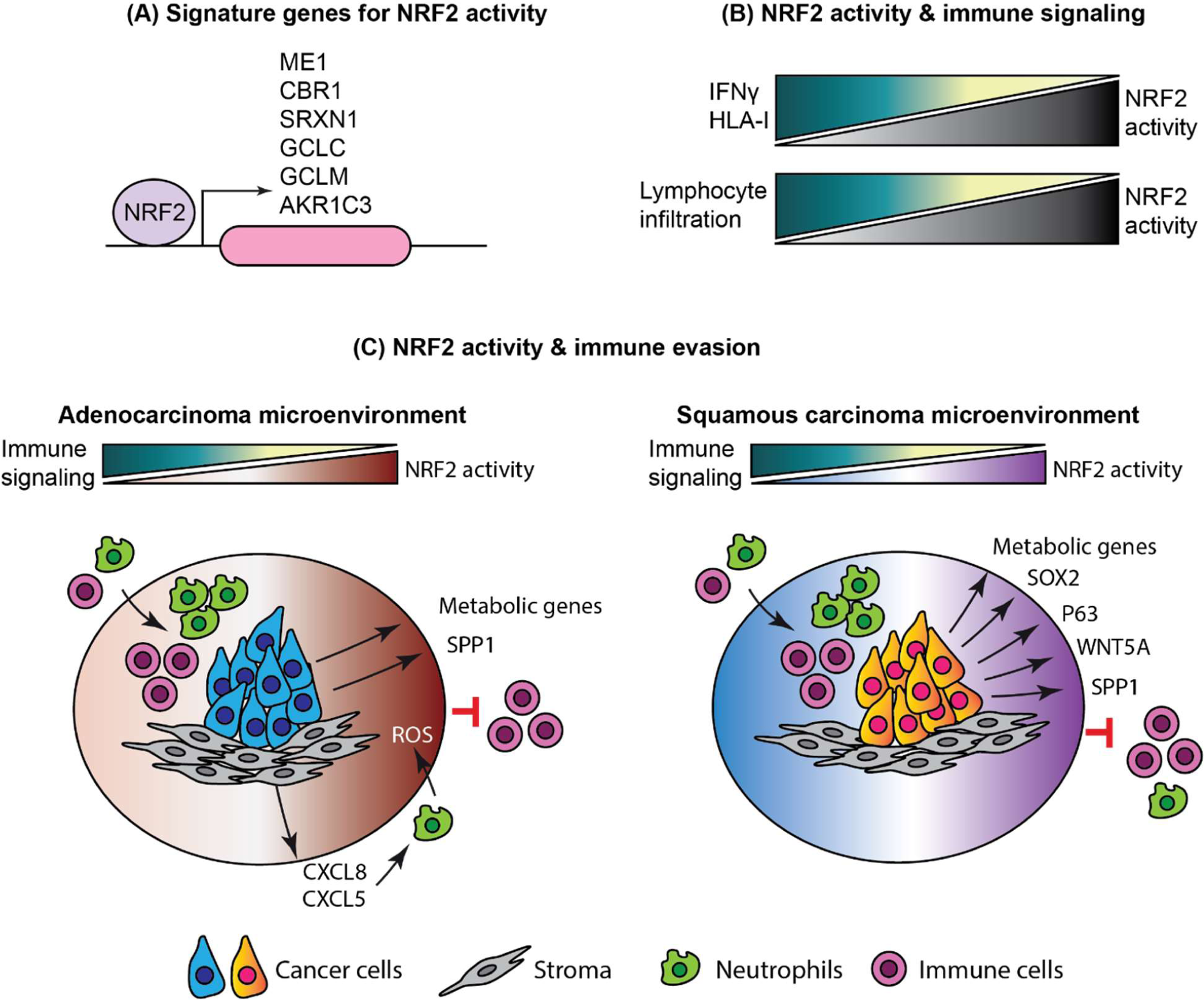

## Introduction

Carcinogenesis – the gradual path from normal cellular behavior to malignant growth – requires a cell to acquire a set of distinct hallmark properties over time. These properties comprise functional changes in pre-malignant cells, as well as in various proximate stromal cells through intercellular crosstalk. From this standpoint, malignant tumors can be considered as complex organs harnessed to sustain neoplastic growth (Hanahan and Weinberg, 2011). Due to the increase in proliferation and cellular metabolism, an inevitable consequence of malignant growth is increased oxidative stress, which renders pathways with antioxidant effects under positive selection (Hayes, Dinkova-Kostova and Tew, 2020). The principal regulator of the cellular redox homeostasis is the transcription factor Nuclear factor erythroid 2-related factor 2 (NRF2, *NFE2L2* gene), which is frequently hyperactive in malignant disease. Especially in non-small cell lung cancer, the oncogenic activation of NRF2 is highly frequent: somatic NRF2 activating mutations alone are among the most frequently occurring subtype specific aberrations (Campbell *et al*., 2016). The cytoplasmic inhibitor of NRF2, Kelch-like ECH Associated Protein 1 (KEAP1), is an E3 ubiquitin ligase substrate adaptor targeting NRF2 for proteasomal degradation under unstressed conditions. In oxidative or electrophile stress, the interaction is disrupted and *de novo* synthesized NRF2 is translocated to the nucleus to drive target gene expression. NRF2 target genes have antioxidant and detoxifying effects via various mechanisms including upregulation of glutathione S-transferases, as well as NAD(P)H quinone oxidoreductase, which has multiple roles in adaptive cellular responses to stress (Ross & Siegel, 2017). In cancer, the regulation of NRF2 is disturbed rendering NRF2 constitutively active. Mechanisms of NRF2 activation include somatic mutations and copy-number variation in *NFE2L2* (gain-of-function or amplification) and *KEAP1* (loss-of-function or deletion), as well as positive regulation by p62 (Leinonen *et al*., 2014; Pölönen *et al*., 2019). Along with its antioxidant effects, NRF2 hyperactivity is known to promote cancer cell proliferation and survival via various other mechanisms, e.g. by promoting anabolic metabolism and increasing chemoresistance via enhanced phase II enzyme and drug efflux transporter expression (Rojo de la Vega, Chapman and Zhang, 2018).

One hallmark property of cancer is its ability to evade destruction by the immune system (Hanahan and Weinberg, 2011). The past decade has complemented oncogene-centric targeted therapies with treatments that modulate the antitumor immune response. However, current diagnostic approaches fail to detect the clinical responders for immune-checkpoint blockade (ICB) with reproducible precision (Ancevski Hunter et al., 2018). From this vantage, the effect of frequent oncogenic events on the crosstalk between cancer cells and immune cells needs further elucidation to dissect the contribution of major oncogenic processes on tissue microenvironment (TME) modification and to provide novel predictive biomarkers for ICB. It has been previously shown that NRF2 drives PD-L1 expression and leads to reduced leukocyte infiltration in mouse allograft models of melanoma and lung adenocarcinoma (Best *et al*., 2018; Zhu *et al*., 2018), but a comprehensive characterization of the effect of NRF2 overactivity on cancer immunity is currently lacking. The aim of this study was therefore to characterize NRF2 hyperactivity at a pan-cancer scale and interrogate unknown biological effects of oncogenic NRF2 activation, with a special emphasis on cancer immunity. To this end, we utilized The Cancer Genome Atlas (TCGA) and Cancer Cell Line Encyclopedia (CCLE), which are publicly available multi-omics databases of tumors and cell lines, respectively. Furthermore, the key findings of this work were validated experimentally and by using independent clinical cohorts.

## Results

### NRF2 activity score improves detection of NRF2 driven malignancies

We developed a NRF2 activity scoring metric from experimentally confirmed robust NRF2 target genes (Figure 1A). The utility of such a score is to overcome the limitations of using gold-standard mutations as a classifier, namely low sample size and the lack of statistical power, as well as ambivalent functionality of rare variants. Furthermore, alternative NRF2 activation mechanisms exist (Leinonen *et al*., 2014; Pölönen *et al*., 2019), highlighting the importance of using NRF2 target gene expression as a marker of activity instead of somatic variants. Our goal was to generate a metric that is: a) based on expression of evident target genes with robust responses to NRF2 activity; b) tissue-agnostic; and c) unbiased towards signal arising from the TME. The score was developed as follows: first, we used A549 lung adenocarcinoma cells harboring the KEAP1 inactivating mutation G333W rendering NRF2 overactive (A549-NRF2^OE^) and knocked out NRF2 with Cas9-sgNRF2 (comparison of A549 NRF2 overexpressed vs knockout, hereafter referred to as A549-NRF2^OEvsKO^) to detect NRF2 dependent differentially expressed genes (Figure S1A & S1B, list of genes in supplementary table 1). Second, we used publicly available functional genomics data (See materials & methods, *Functional genomics analysis of NRF2 target genes*) to subset the genes that are directly regulated by NRF2 (See supplementary table 2 for a complete reference of candidate targets). Third, we utilized public tissue as well as stromal cell transcriptomic data from Genotype Tissue Expression portal (GTEx) and Database of Immune Cell Expression (DICE), respectively, to discard genes that are clearly tissue specific or prominently expressed in TME-populations (Figure S1C and Figure S1D). The final NRF2 signature comprised genes *CBR1, SRXN1, GCLC, GCLM, AKR1C3* and *ME1* and the final score was defined as a geometric mean of their linear TMM normalized mRNA-expression. The score values were scaled within disease entities (to density distribution peak) in TCGA-samples to decrease variance between cancer types.

**Figure 1:**
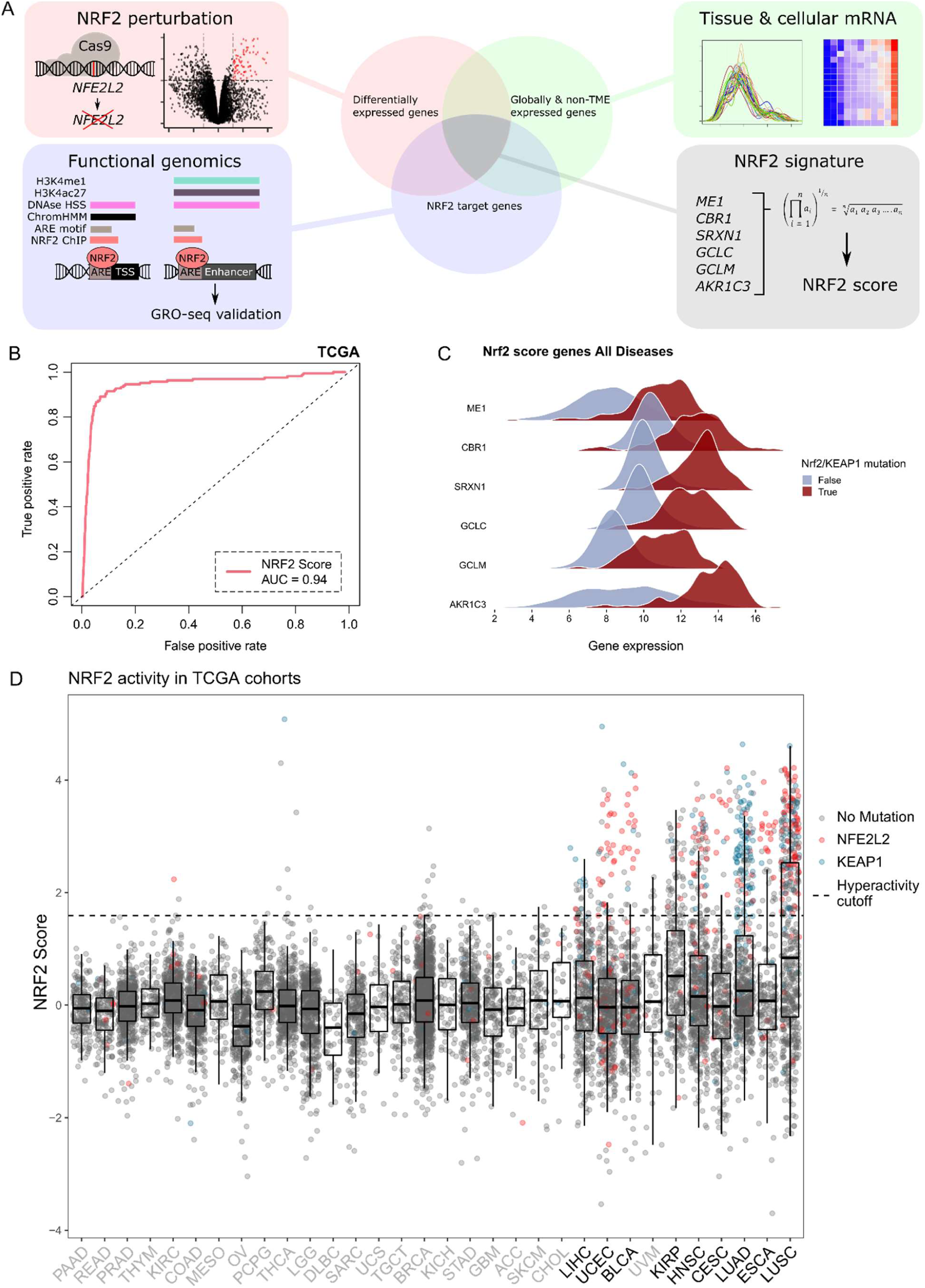
NRF2 score development, performance and distribution across TCGA malignancies. **A:** Schematic of the score development procedure. **B:** Receiver operating characteristic (ROC) curve for the NRF2 score in classifying functionally relevant somatic variants defined in OncoKB. **C:** Differential log2 CPM gene expression distribution between *NFE2L2/KEAP1* mutated and non-mutated NRF2 score comprising genes in TCGA cohorts. **D:** NRF2 score with *NFE2L2/KEAP1* mutations shown as a boxplot across TCGA-cohorts sorted by variance. The cohorts chosen into subsequent analyses are colored in black. The hyperactivity threshold was chosen with values of TPR > 0.85 and FPR < 0.1.

We assessed the score performance with ROC-analysis against OncoKB (Chakravarty *et al*., 2017) defined hotspots in *NFE2L2* and *KEAP1* and functional variants (*KEAP1*-truncating aberrations) in TCGA-data (Figure 1B). The score exhibited excellent overall discrimination (AUC = 0.94) and all of the genes showed markedly different distributions in mRNA-expression in TCGA *KEAP1*/*NFE2L2* mutated vs wild type samples (Figure 1C). Individual ROC analyses for cohorts with somatic mutations in *NFE2L2*/*KEAP1* are shown in Figure S1E. Score variance correlated considerably with *KEAP1*/*NFE2L2* mutation frequency (r = 0.8, P < 0.0001) and thus evidently predicts the oncogenicity of NRF2 irrespective of activating mechanism (Figure S1F). By this rationale, diseases associated with > 5 % mutation frequency or σ^2^ > 0.75 were defined to harbor significant oncogenic NRF2 activity. From these malignancies, we discarded TCGA cohorts with N < 100 to maintain high statistical power across the datasets. The score distribution and cohort selection is shown in Figure 1D (colored mutations above the specified hyperactivity cutoff of TPR > 0.85 are shown in Figure S1G. NRF2 activity scores for all TCGA samples are listed in supplementary table 3). Notably, malignancies in the lung, uterus, bladder, kidney and those with squamous histology had a significant proportion of high scoring samples.

### Enriched pathways in NRF2 overactive cancers reveal differential immunomodulatory association across cancer types

For TCGA cohorts meeting the inclusion criteria, as well as for the A549-NRF2^OEvsKO^ and CCLE transcriptome and proteome data, we conducted gene-set enrichment analysis (GSEA) to assess the pathways enriched with NRF2 hyperactivity (Figure 2A. See supplementary table 4 for all data). Interestingly, two directly oncogenic signaling pathways, MYC and WNT, showed prominent global positive enrichment. Other global pathways associated with NRF2 hyperactivity were mainly metabolic, drug efflux and redox-regulatory processes, whereas immune microenvironment related processes were negatively enriched in NSCLC and squamous diseases in contrast to other diseases, which exhibited positive enrichment. With the curated data, cohorts clustered into two populations based on the immune milieu associated gene sets. Notably, IFNγ response, HLA-and T-cell signaling gene sets enriched to the negative end in lung adenocarcinoma (LUAD), lung squamous cell carcinoma (LUSC), esophageal carcinoma (ESCA), cervical carcinoma (CESC) as well as head and neck cancer (HNSC), while the same pathways had positive enrichment scores in kidney renal papillary cell carcinoma (KIRP), uterine corpus endometrial carcinoma (UCEC), bladder carcinoma (BLCA) and liver hepatocellular carcinoma (LIHC) (Figure 2A and 2B). As the immunological gene-sets were not enriched in the pure cell populations (A549-NRF2^OEvsKO^ or CCLE), they likely emerge from the crosstalk between cancer-and TME-cell populations. Of note, downregulation of cytokines, HLA-I, IL-12, IFNγ and TCR signaling are all characteristic to ‘immune cold’ tumors with documented poor responses to ICB therapies (Duan *et al*., 2020). Upon further characterization of the genes responsible for the negative enrichment in the TME-associated pathways, we observed genes linked to both major HLA-I and HLA-II associated antigen presentation as well as T-cell co-receptors in the antigen processing and presentation pathway gene set (Figure 2c). Since HLA-II genes are expressed only in professional antigen presenting cells and *CD4, CD8A* and *CD8B* are exclusive to T-cells, whereas HLA-I genes are also expressed in cancer cells, these data suggest that oncogenic NRF2 signaling is, in the context of NSCLC and squamous diseases, associated with less lymphocyte infiltration and/or a weaker response to IFNγ. The data also shows a dichotomous association of NRF2 to IFNγ high and low tumors.

**Figure 2:**
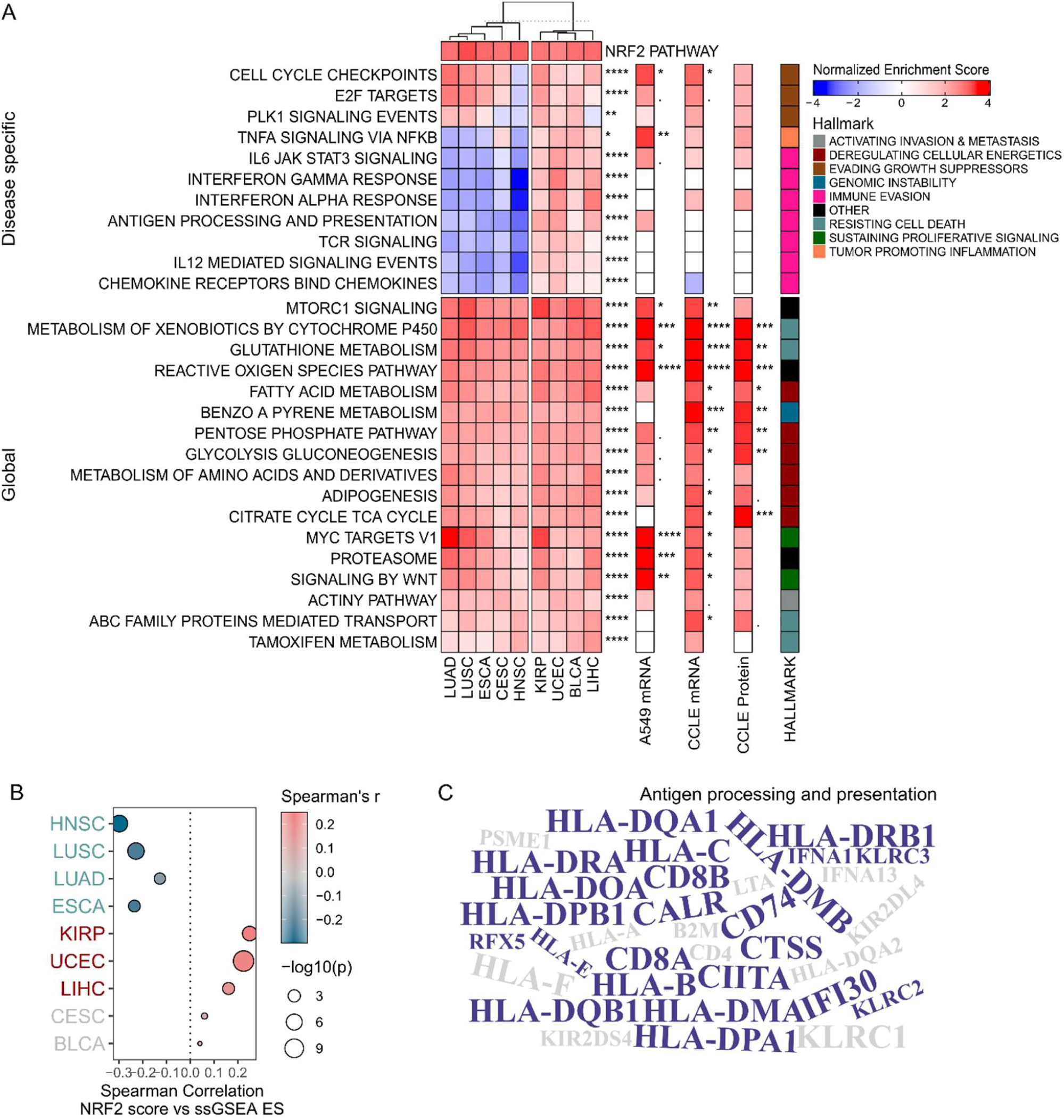
NRF2 activity associates with disease specific immunoevasive characteristics. **A:** Pan-cancer GSEA analysis. GSEA normalized enrichment scores for pathways correlated to NRF2 activity shown as a Heatmap for TCGA, A549-NRF2^OEvsKO^, CCLE mRNA and CCLE protein. Significance of enrichment is shown as: < 0.1, * < 0.05, ** < 0.01, *** < 0.001, **** < 0.0001. For TCGA, combined p-values (Stouffer’s method) are shown. Redundant pathway terms were discarded, See Table S2 for full list of pathway terms. Pathways were curated to reflect cancer hallmarks that are shown on the right. **B:** Correlation of ssGSEA Hallmarks IFNγ response and NRF2 score. Cohorts in blue, red and grey have negative, positive or insignificant correlation, respectively. **C:** Word cloud of the core-enrichment terms in the antigen processing and presentation pathway within the negatively correlating cohorts.

### Squamous diseases comprise a distinct NRF2 hyperactive subtype with differentially expressed metabolic genes inflicting resistance to T-cell mediated killing

Since we observed similar immune-cold characteristics across the TCGA squamous diseases, we proceeded to study subtype-effects within the diseases using Uniform Manifold Approximation and Projection (UMAP) and community detection-based clustering. We identified a distinct pan-squamous subtype with hyperactive NRF2 (identified communities are shown in Figure 3A, TCGA cohorts in Figure 3B and NRF2 activity in Figure 3C). Associated to this subtype, we observed co-occurring copy number variation (CNV) and a characteristic mutational landscape, most notably amplified *SOX2*/*TP63* (q-arm of chromosome 3) and loss of *CDKN2A*/*CDKN2B* (9p21) (Figure 3D and S3 A-D), as well as mutated *TP53* and *CDKN2A* (Figure S3 e & F). Furthermore, we identified a group of cell lines with similar genomic profiles using the CCLE/DepMap dataset, confirming the genomic determinants of this subtype (Figure S3G, H, I, J, K, L, and M). There were no prominent peaks in chromosome 6, suggesting that HLA-I loss events do not contribute to the phenotype. To confirm this, we defined HLA-I loss as at least one shallow deletion or LoF-mutation in major HLA-I genes, and did not observe an association between the two variables with Fisher’s test (OR = 0.40, P = 0.09). The association of tumor mutational burden and NRF2 overexpression has been reported before (Liu *et al*., 2021). Thus, we assessed the prospect of immunological effects arising from differential neoantigen load by computing mutational burden (log2 total mutation count) across the cluster comprising cohorts with respect to NRF2 activity, and did not observe a uniform trend between mutation count and NRF2 (Figure S3N). To follow up on amplified transcription factors *SOX2* and *TP63*, we downloaded publicly available ChIP-seq data (GSE46837) in squamous cancer cell lines and generated a list of target genes (pipeline as in Figure 1A, functional genomics) to interrogate direct targets of these transcription factors (see supplementary tables 5 and 6 for a complete reference of targets). To specifically assess differentially expressed, putative direct targets in immunomodulatory genes, we studied the overlap of the differentially expressed genes in the squamous cluster and the target gene list as well as curated genes from the CellPhoneDB framework (Efremova *et al*., 2020) and TISIDB (Ru *et al*., 2019) (Figure 6E). From these results, *SLC7A11, NAMPT, SLC2A1* and *MPP3* were identified to confer resistance to T-cell mediated killing by high-throughput screening. Moreover, all of these genes had evidence for upstream activation by NRF2. *SLC7A11, NAMPT* and *SCL2A1* are genes attributed to metabolism, involved in cysteine uptake, NAD+ biosynthesis and glucose uptake, respectively. From the literature annotated genes, while most were ambivalent, *SOX2* was shown to be a) upregulated in the case of effector cell resistance in co-culture; b) downregulate IFN type I response *in vitro* and c) decrease T-cell infiltration in a HNSC murine model (Tan *et al*., 2018). Interestingly, we observed increased *NFE2L2* mRNA in *SOX2* amplified TCGA squamous cell carcinoma (SqCC) cases, suggesting the presence of an upstream regulator in Chr3 q2 locus (Figure S3O). Finally, in the squamous NRF2 cluster, we identified prominent downregulation of the IFNγ response similar to the initial GSEA analysis (Figure 3F). Taken together, if the immunological effects are attributable to oncogenic signaling, the downregulation of IFNγ-response and/or HLA-I genes may be downstream of NRF2 or other co-expressed transcription factors (SOX2 or TP63). Moreover, based on the identified hits with functional evidence (Figure 3E), metabolic reprogramming and the metabolic microenvironment may prove relevant to the immunoevasive characteristics of NRF2 driven malignancies.

**Figure 3:**
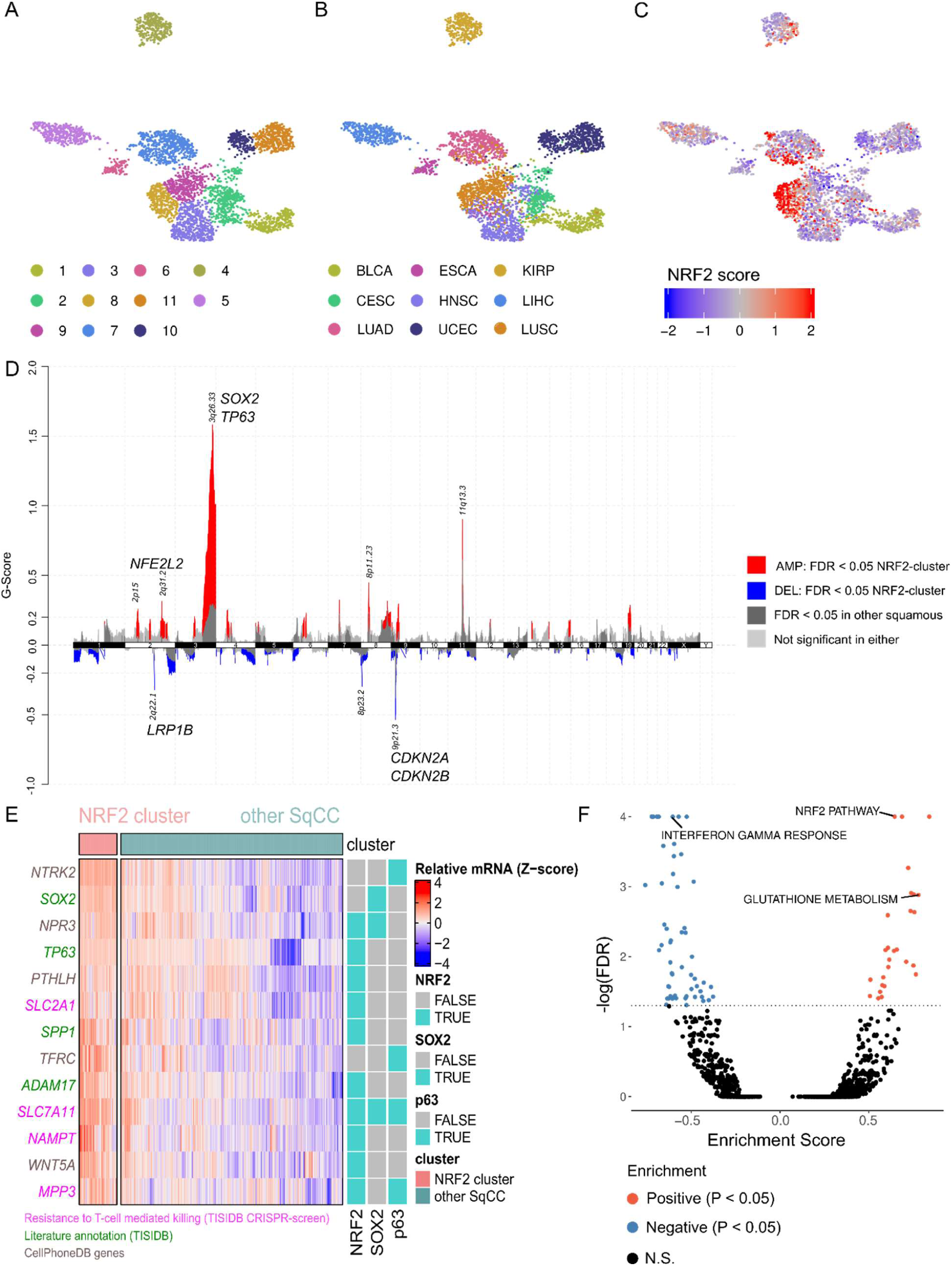
NRF2 hyperactivity is associated with a distinct squamous subtype with *SOX2/TP63* amplification, *CDKN2A/CDKN2B* loss and downregulated IFNγ-response in TCGA. **A:** UMAP representation and Louvain clustering of included TCGA cohorts. **B:** TCGA diseases in UMAP clusters. **C:** NRF2 score in UMAP clusters. **D:** Gistic2 copynumber profile of the NRF2 hyperactive cluster*. The G-score in gistic2 is CNV amplitude multiplied by CNV frequency, which measures the significance of a CNV event in a dataset. **E:** Differentially expressed immunomodulatory genes between the NRF2 hyperactive cluster and other SqCC putatively regulated by NRF2, SOX2 or TP63. From the TISIDB-database, genes colored with green indicate a literature cited effect, whereas genes colored in magenta depict a hit from CRISPR/Cas9 functional screens. **F:** Enriched pathways in the NRF2 hyperactive cluster*. *All analyses were conducted against squamous samples; LUSC, HNSC, ESCA and CESC.

### Interferon-gamma response pathway is downregulated in NRF2 hyperactive cancer cells *in situ*

To follow up on the identified targets, we proceeded to further explore signaling between NRF2 hyperactive cancer cells and immune cells in higher resolution in an *in situ* setting in a relevant cancer type. Thereby, we assigned NRF2 activity score to cells in a publicly available HNSC single-cell-RNAseq dataset (GSE103322). UMAP projection of single-cells revealed distinct clustering of cancer cells with high NRF2 score across patient samples, suggesting that NRF2 activation could be linked to global shifts in cell phenotype (Figure 4A and 4B). Similar to bulk tumors, the cluster exhibited high expression of *SOX2* (Figure 4C). In addition, *TP63* was also overexpressed in the NRF2 hyperactive cancer cell cluster, although its expression was also present in other clusters (Figure S4A). In further agreement with the bulk-tumor analysis, inflammatory response- and IFNγ-signaling were the most prominent negatively enriched pathways in the NRF2 cluster relative to other cancer cell clusters (Figure 4D). The single cell analysis distinguished that the response to interferon is downregulated in malignant cells with NRF2 hyperactivity (Figure S4B). These data suggest that the negative correlation between IFNγ response and NRF2 in squamous cancer types originates from the response in cancer cells, either due to low interferon ligand or by intrinsic properties of NRF2 hyperactive cancer cells.

**Figure 4:**
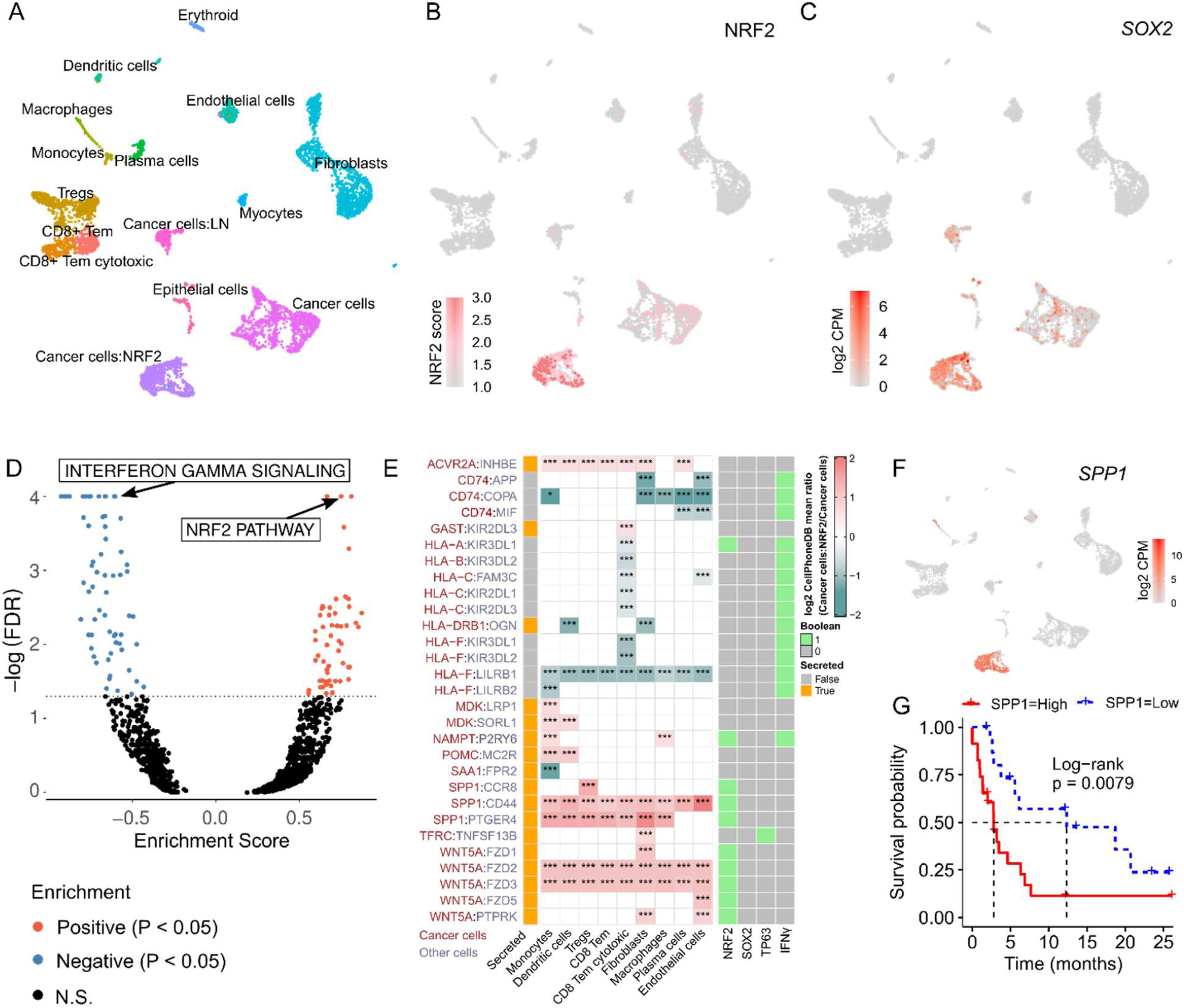
NRF2 hyperactive cancer cells have a dampened IFNγ response *in situ* and exhibit unique immunomodulatory intracellular interactions. **A:** UMAP representation for cell populations. **B:** UMAP representation for NRF2 score. **C:** UMAP representation for *SOX2* mRNA. **D:** GSEA of NRF2 hyperactive vs other cancer cells. **E:** Curated interactions from the CellPhoneDB analysis. Statistical significance denoted as: * P < 0.05; * P <0.001. **F:** UMAP representation for *SPP1* mRNA. **G:** Kaplan-Meier survival for *SPP1* high vs low in pan-HNSC-LUAD-LUSC.

### NRF2 hyperactive cancer cells are associated with less TME interactions via HLA-I and increased interactions via *NAMPT, SPP1* and *WNT5A*

We used the statistical framework of CellPhoneDB to interrogate putative intercellular ligand-receptor interactions between cancer- and TME cells in the whole HNSC single-cell dataset. With integration of our NRF2 target catalogue and a priori IFNγ gene sets from MSigDB (Hallmarks and Reactome), we identified multiple differential ligand-receptor interactions between the cancer clusters against TME-clusters with either direct NRF2 targets or genes involved in IFNγ mediated signaling. The most prominent hits were downregulated HLA type I interactions with cytotoxic T-cells, and upregulated *NAMPT, SPP1, WNT5A* (Figure 4E and 4F), all of which were also differentially expressed genes in the earlier pan-SqCC bulk tumor analysis and in CCLE cell lines mRNA, while *SPP1* and *WNT5A* proteins were also upregulated (Figure S4C, D and E). Furthermore, *NAMPT* and *SPP1* were differentially expressed in A549^OEvsKO^ and all of the hits were in our NRF2 target catalogue (Figure S4F). Moreover, in line with earlier observations, PD-L1 -PD1 interaction between NRF2 hyperactive cancer cells and T-cells was statistically significant, further corroborating the role of PD-L1 in NRF2 driven immune-escape, and suggesting that its effect extends the previously studied melanoma (Zhu *et al*., 2018) (Fig S4G). From the direct targets, NAMPT is the rate-limiting enzyme in the biosynthesis of NAD+. While its interaction with P2RY6 in CellPhoneDB was inferred with protein pulldown and lacks functional data, NAMPT knockdown was recently shown to increase CD8+ T-cell infiltration in murine tumors via attenuating inducible PD-L1 expression (Lv *et al*., 2021). The second hit, Osteopontin (*SPP1*), has been shown to inhibit T-cell proliferation and IFNγ production *in vitro* (Klement *et al*., 2018). The third hit, WNT5A with frizzled receptors, also has implications in tumor immunity: WNT5a signaling in cancer cells has been attributed to immunosuppressive metabolite induction via dendritic cells through FZD (Zhao *et al*., 2018). To assess the clinical relevance of these findings, we downloaded a publicly available targeted mRNA-expression dataset of PD-1 inhibitor treated HNSC and NSCLC patients and established *SPP1* mRNA as a negative predictive biomarker to treatment-response (Odds ratio = 0.20 for response with high *SPP1* expression; P < 0.05, Fisher’s test) and therapy associated progression-free survival (mPFS 2.8 vs mPFS 12.33 months, P < 0.01) in pan HNSC-LUSC-LUAD (n = 5; n = 22; n = 13, respectively) (Figure 5G). The inferior survival may either be directly caused by effects of SPP1, or arise from NRF2 activation where *SPP1* mRNA acts as a surrogate marker. From a biomarker perspective, the clinical observation is relevant for classifying non-responders to ICB, since currently the insight on ICB treatment response in cancer patients is widely elusive. In summary, these data demonstrate that NRF2 hyperactive cancer cells overexpress unique TME interacting ligands *NAMPT, SPP1* and *WNT5A*, which have been shown to cause immunoevasion in cancer.

**Figure 5:**
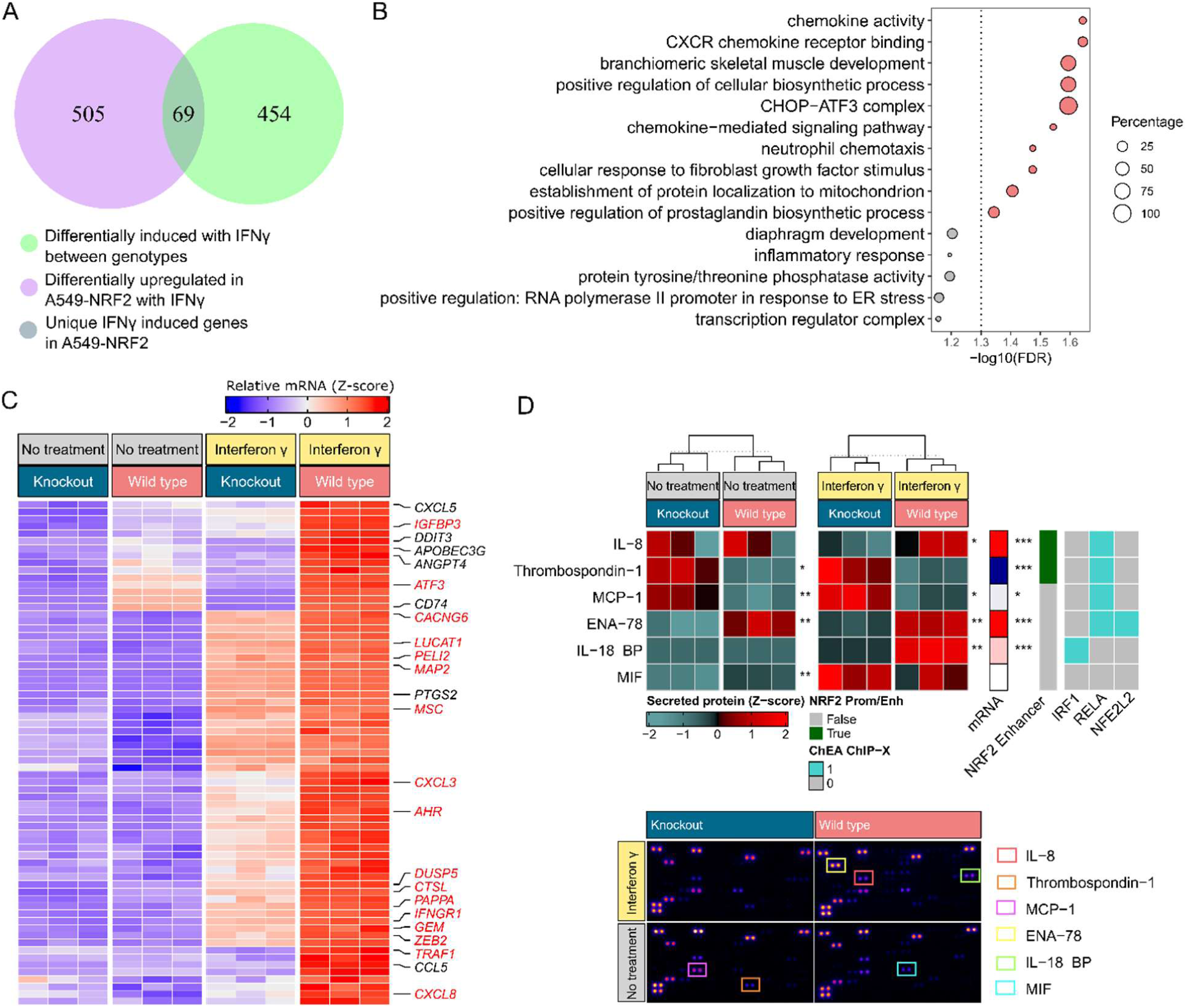
IFNγ differentially induces immunomodulatory genes and cytokine secretion in A549-NRF2^OEvsKO^. **A:** Venn-diagram of genes overexpressed in A549-NRF2 and genes differentially induced between genotypes with IFNγ. **B:** Gene set enrichment analysis (GeneSCF) for the IFNγ induced genes overexpressed in A549-NRF2^OEvsKO^ with IFNγ treatment. **C:** Heatmap of IFNγ induced genes overexpressed in A549-NRF2^OEvsKO^ with IFNγ treatment. Immunogenomic terms and putative NRF2 targets (colored red) are annotated on the right. **D:** Proteome profiler for human cytokines in A549-NRF2^OEvsKO^ ± IFNγ.

### NRF2 regulates IFNγ-responsive genes in lung adenocarcinoma *in vitro*

The single-cell data indicated that the repression of IFNγ-responsive genes in the NRF2 hyperactive tumors may be caused either by low production of IFNγ ligand (that is, negligible infiltration of proinflammatory cells) or is attributed to intrinsic properties of the NRF2 hyperactive cancer cells (lower response to IFNγ). IFNγ binding to its receptor cause a cascade of phosphorylation events which activates the transcription of interferon-stimulated genes via phosphorylated nuclear localized STAT1. Among the plethora of its effects, IFNγ stimulation leads to direct effector functions in many immune cell types, and to the expression of MHC I genes for neoantigen presentation in cancer cells (Ivashkiv, 2018). We sought to assess the causality by subjecting the A549-NRF2^OEvsKO^ cells to IFNγ and measuring the transcriptomic response with RNA-seq. The IFNγ pathway was not downregulated by NRF2 in response to IFNγ (Fisher’s test, OR = 1.26, P = 0.37, Hallmarks gene set). However, IFNγ treatment revealed a distinct set of differentially expressed genes, potentiated or exclusive to IFNγ treatment, between the genotypes (Figure 5A and Figure S5A). Gene set enrichment analysis (performed with GeneSCF, Subhash and Kanduri, 2016) revealed enriched processes within these genes, most notably increased chemokine expression and neutrophil chemotaxis (positive end in Figure 5B). To characterize transcriptional changes attributable to immunological processes and NRF2 transcriptional activity, we integrated the IFNγ induced differentially overexpressed genes with our NRF2 target catalogue as well as genes defined in TISIDB and InnateDB (Breuer *et al*., 2013) (Figure 5C). Based on our NRF2 target list, 25 % of these genes had evidence for promoter or enhancer mediated regulation via NRF2. The main analyses were conducted for the upregulated genes due to higher amount of hits and context relevance of the associated pathway terms (downregulated genes are presented in Figure S5A-C). To assess other putative transcriptional events responsible for the differentially overexpressed genes in the IFNγ treatment group, we utilized the ChEA Transcription Factor Targets (ChEA-TF) dataset (Lachmann *et al*., 2010) for the assessment of overrepresented transcription factors in the gene list using Fisher’s test. The most prominently enriched transcription factors in addition to ARE-binding NRF2 and BACH1, excluding explicit repressors, were RELA and IRF1 (Figure S5D). Based on the ChEA-TF dataset and our NRF2 enhancer catalogue, RELA, IRF1 and NRF2 together covered 68 % of the A549-NRF2^OEvsKO^ differentially upregulated genes in the IFNγ treatment group (Figure S5E). Due to the clear difference in expression patterns of several cytokines (*CXCL8, CXCL5, CXCL3* and *CCL5*), we went ahead to explore paracrine immunomodulatory signaling arising from the differential response to IFNγ. To this end, we collected conditioned media (CM) from the cells with and without IFNγ and utilized a cytokine array for detection of 104 targets. Most notably, we observed increased secreted IL-18 binding protein (IL18BP) and IL-8 (*CXCL8*) in the A549-NRF2^OEvsKO^ conditioned media with IFNγ-treatment (Figure 5D). In adfdition, ENA-78 (*CXCL5*) and MIF were increased regardless of treatment and in the untreated group, respectively, while Thrombospondin-1 (*THBS1*) and MCP-1 (*CCL2*) were downregulated in the untreated group (Figure 5D). IL18BP is a potent antagonist of IL-18. IL-18 stimulates multiple immune cell types to produce IFNγ and induces effector functions in T-cells (Zhou *et al*., 2020). ENA-78 and IL-8 are both neutrophil chemoattractants that mobilize neutrophils to the tumor site (Wu *et al*., 2019). Based on the ChEA-TF analysis and NRF2 enhancer catalogue, *IL18BP* had an IRF1 peak on its promoter, while *CCL2* (MCP-1) had RELA, and *CXCL8* (IL-8) as well as *CXCL5* (ENA-78) had peaks for NFE2L2 and RELA. In this particular model, there was no difference in secreted *SPP1*, while its mRNA was differentially expressed (logFC = 2.6; FDR < 0.0001). While this data suggests that the negatively correlating IFNγ response in tumors is not attributable to NRF2-dependent cancer intrinsic attenuation of the IFNγ mediated signaling, it shows that the response in a subset of IFNγ responsive genes is elevated in NRF2 hyperactive cells, connecting neutrophil mediated immune suppression and inhibition of T-cell effector phenotypes via IL18BP with NRF2 biology in the context of cancer.

### NRF2 hyperactivity associates with reduced total tumor lymphocyte infiltration in NSCLC and pan-squamous cancers

Finally, to explore the relationship between NRF2 activity and the immune-milieu in greater detail, we performed deconvolution for the TCGA gene expression data with CIBERSORT (Newman *et al*., 2015) to infer immune cell content, and correlated the inferred cell type fractions to the NRF2 score (Figure 6A). In the NRF2 hyperactive IFNγ negative malignancies (cohorts color coded as cadet blue), many lymphocyte populations and different macrophage polarization states correlated negatively with NRF2 activity, while the correlations of cell subsets varied between cancer types. In the NRF2 hyperactive IFNγ positive cancers (cohorts color coded as coral), we observed mostly positive correlations to different immune cells with variability in the correlating cell types between diseases: in KIRP, there was a strong association to macrophages and in UCEC to other antigen presenting cells. Due to high underlying variability in the immune landscape between different malignancies (Thorsson *et al*., 2018), we performed Mann-Whitney U tests for the immune cell content between the IFNγ response curated cancers, and observed that the negatively correlating diseases often had a higher disease-intrinsic content of the assessed cell types (Figure 6A, side panel). To evaluate the relationship of NRF2 and total T-lymphocyte content, we calculated the sum of T-cell fractions and observed less bulk lymphocytes in all of the IFNγ-negative NRF2 hyperactive cancers (Figure 6B). We proceeded to validate the result in LUSC with a different approach utilizing a publicly available TIL dataset generated from TCGA H&E images with deep learning (Saltz *et al*., 2018), and observed a decrease in the median percentage of tumor-infiltrating lymphocytes (TILs) in the NRF2 hyperactive cases (P < 0.05, Figure 6C). To corroborate this, we stained an independent tissue-microarray dataset of NSCLC (117 and 211 cases of LUSC and LUAD, respectively) with NQO1 (a marker of NRF2 activity) and the universal T-cell marker CD3, and observed a similar result in LUSC, that is, NQO1 positive cases harbored less intratumoral CD3-positive cells (P < 0.05, Figure 6D-E). In our TMA-data, the cytotoxic T-cell marker CD8 showed a similar trend, but did not reach statistical significance (P = 0.09, Figure S6A), and in LUAD, the relationship was not observed (Data not shown). In addition, the contribution of *NFE2L2* and *KEAP1* mutations was assessed separately in TCGA LUSC cohort, and a lower median fraction of total lymphocyte infiltration was present in all groups, irrespective of mutated gene (Figure S6B). Taken together these data suggest that overactive NRF2 associates with less overall lymphocyte infiltration in NSCLC and pan squamous cell carcinoma, with variations in the specific subtypes of reduced lymphocytes between cancers.

**Figure 6:**
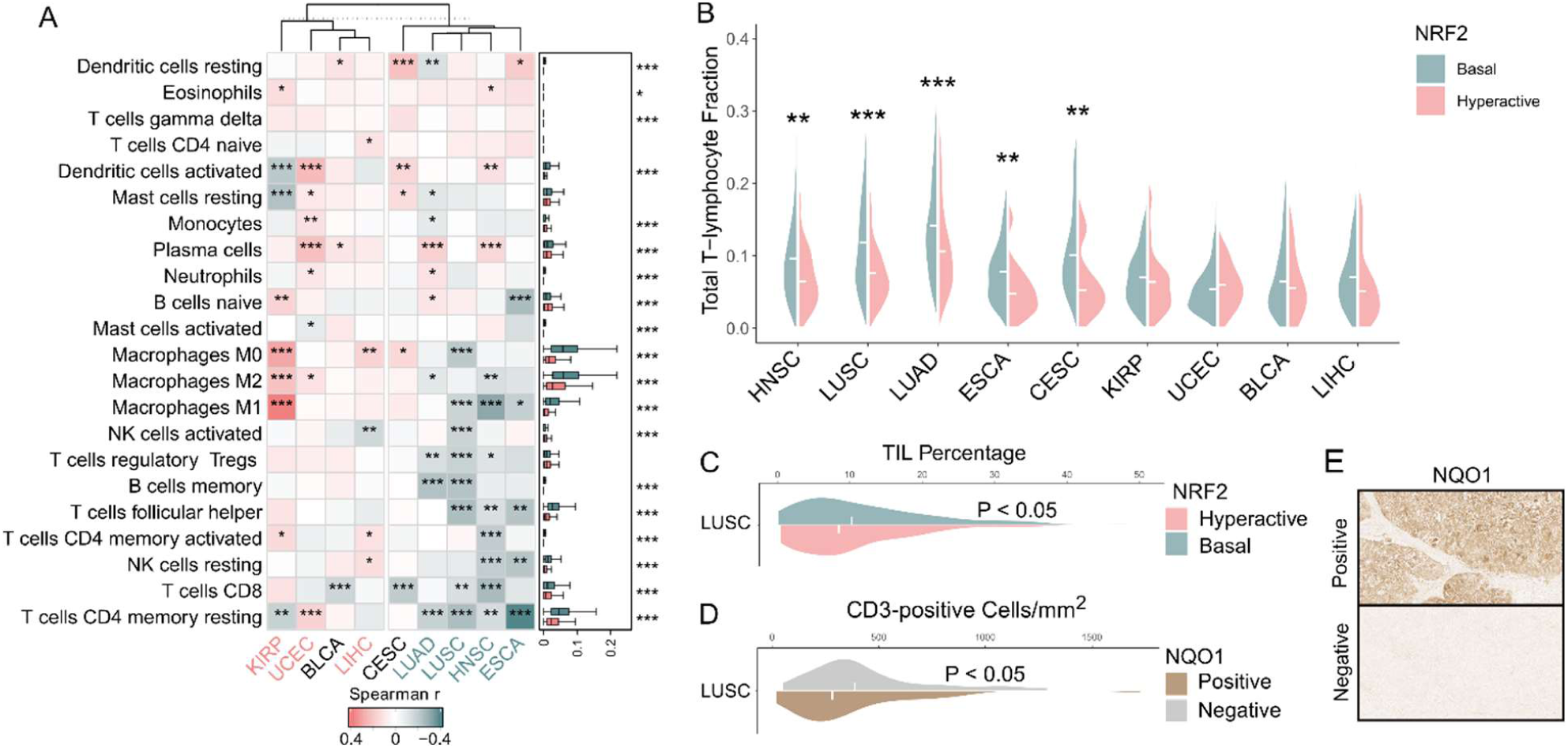
NRF2 hyperactive tumors display negative correlations to overall lymphocytes in NSCLC and pan-squamous diseases. **A:** Correlation matrix of NRF2 score vs CIBERSORT cell types. Right annotation panel depicts the distributions of respective immune cells in the IFNγ pathway classified diseases. Statistical significance is shown as: * P < 0.05, ** P < 0.01, *** P < 0.001. **B:** Violin plots for CIBERSORT bulk T-lymphocyte fractions. **C:** Percentage of tumor-infiltrating lymphocytes (TIL) in digitized TCGA LUSC H&E images. **D:** Tissue microarray IHC staining of NQO1 (NRF2 activity marker) and CD3+ cells. **E:** Representative images of the NQO1-stained groups.

## Discussion

Cancer immunotherapies, especially ICB, has become a mainstay in cancer care. Despite its undeniable success, biomarkers to stratify responders and non-responders are currently elusive. Recently, the relationship between oncogenic pathways and the TME have been explored. For instance, the effect of MYC and WNT signaling, as well as the loss of TP53, LKB1 and PTEN on tumor immune responses have been previously characterized (Spranger and Gajewski, 2018). In this work, we identified a phenotype of high NRF2 activity accompanied with low lymphocyte content, HLA-I and IFNγ, spanning multiple cancer types. Since the negative relationship between immune infiltration and immunotherapy responses has been clearly established before, it is reasonable to assume that NRF2 hyperactive cases of the characterized malignancies would associate with weaker responses, and that alternative strategies to activate the antitumor immune response should be studied. Indeed, during the course of this study, an association between NRF2 pathway mutations and inferior ICB-response was shown in NSCLC (Singh *et al*., 2021). The present study, to our understanding, is the first attempt to provide pan-cancer wide perspective to NRF2 signaling and tumor immunity, with important implications for future studies: first, immune checkpoint inhibitor treatment efficacy should be assessed prospectively across all squamous malignancies with NRF2 hyperactivity and eligibility for ICB; and second, if proven inferior, alternative first-line or effective adjuvant treatment options should be explored.

Characterizing cancers by mutational profiling of large datasets has been invaluable in gaining insight into the biological processes enabling and governing malignant growth. However, profiling only by mutational status has limitations: The functional consequences of individual mutations may be ambivalent, other mechanisms for perturbation of the same pathway are overlooked, or the frequency of events are too low for robust statistical inference. Relevant to NRF2 mediated oncogenic signaling, mutation-independent means of activation are common, necessitating the use of gene expression classifier to deduct NRF2 activation (Kansanen *et al*., 2013). Herein, we introduce a robust NRF2 scoring metric comprising genes that have minimal tissue specificity and sound evidence for direct, prominent activation. The score performs well in all of the tested TCGA cohorts (Figure S1E) and it can be applied to bulk tissue and single cell data. Moreover, the metric was constructed to support ease of use by utilizing a low number of genes and prioritizing global, robust targets. Finally, we demonstrated the utility of the score by predicting a distinct immunological phenotype in NRF2 hyperactive cases in an external dataset using IHC staining as an alternative method for NRF2 activity assessment.

Our study provided a number of leads towards elucidating the mechanisms by which NRF2 overactivity affects the host immune response. We outruled the contribution of HLA-I loss and neoantigen effects, and explored IFNγ induced signaling in a lung adenocarcinoma model. We identified a distinct squamous cell subtype that was characterized by NRF2 overactivation together with *SOX2*/*TP63* amplification and immune cold TME. Sox2 is a key driver of malignant transformation and stemness in squamous type cancers (Porter and McCaughan, 2020). Our analysis showed the metabolic genes *SLC2A1, SLC7A11* and *NAMPT* that associate with functional evidence in driving resistance to T-cell mediated killing as downstream candidates of NRF2. Previous work shows that the expression of *SLC2A1* - a glucose transporter - is enhanced by p63 and Sox2 (Hsieh *et al*., 2019). Our work implicates functional synergy of NRF2 and p63/Sox2 in *SLC2A1* driven glucose uptake. Increased glucose uptake via *SLC2A1* has been shown to result in weaker anti tumoural immune responses in a murine model (Ottensmeier *et al*., 2016), supporting the functional role of this gene identified by the TISIDB high-throughput CRISPR-screen. *SLC7A11*, a well characterized NRF2 target gene (Hayes and Dinkova-Kostova, 2014), is a glutamate-cystine antiporter that increases intracellular cystine while exporting glutamate. IFNγ has been reported to downregulate the glutamate-cystine antiporter members *SLC7A11* and *SLC3A2*, lowering intracellular cystine resulting in increased T-cell induced ferroptosis (Wang *et al*., 2019). Thus, the higher expression of *SLC7A11* in NRF2 overactive cancers may render the cells resistant to immune destruction by this mechanism. NAMPT driven NAD+ metabolism is involved in tumor immunity: increased NAD+ by NAMPT enables inducible PD-L1 expression by epigenetic regulation of Irf1, restricting the antitumor action of cytotoxic T-cells (Lv *et al*., 2021). The results suggest metabolic crosstalk between NRF2 overexpressing and TME cells through multiple mechanisms.

*SPP1* and *WNT5A* were hits in both the bulk tumor and single cell analysis. Both genes have implications in tumor immunity. In line with the immunoevasive phenotype, soluble *SPP1* has been shown to directly inhibit T cell proliferation, decrease T-cell activation and IFNγ production *in vitro* (Klement *et al*., 2018). The observed interaction of WNT5A and frizzled receptors in the single-cell data has intriguing implications in dendritic cell (DC) function: paracrine WNT5A signaling was shown to induce IDO1 in DCs, which increased extracellular kynurenine, supporting an immunosuppressive TME (Zhao *et al*., 2018).

In lung adenocarcinoma, immune-evasion via neutrophil recruitment and/or inhibition of effector phenotypes via antagonism of IL-18 were avenues revealed by comparing NRF2 overexpressing A549 vs knockout cells. Signaling associated with these processes was clearly different: ENA-78 (*CXCL5*), IL-8 (*CXCL8*) and IL-18 binding protein (*IL18BP*) were upregulated in the transcriptome and the secretome of the NRF2 overexpressing cells. In line with the *in vitro* findings, we observed a statistically significant, albeit weak, positive correlation between NRF2 signaling and CIBERSORT inferred neutrophils in LUAD. Recently, neutrophils have been shown to function as immunosuppressive players in the TME. The effect emerges from a subpopulation of suppressive neutrophils that inhibit T-cell proliferation via deprivation of L-arginine and by generating ROS (Granot, 2019). IL-18 binding protein has been shown to act as a secreted immune checkpoint, contributing to the exhaustion of CD8+ T-cells and inhibition of natural killer cells by inhibiting the IL-18-mediated stimulation and IFNγ secretion of these cells (Zhou *et al*., 2020).

In summary, our results provide an integrated NRF2-centric resource in cancer biology, and establish a connection between immune-cold tumors and NRF2 signaling across NSCLC and squamous carcinomas from a pan-cancer wide perspective. Finally, these data highlight multiple avenues to pursue in future studies aiming to mechanistically characterize NRF2 signaling and its effect to the tumor immune milieu, including direct regulation via immunosuppressive ligands or immunosuppressive metabolites, as well as neutrophil recruitment into the tumor bed.

## Acknowledgements

This study was supported by University of Eastern Finland Doctoral Program in Molecular Medicine, Finnish Cancer Foundation, The Academy of Finland (Grant number 332697), Sigrid Juselius Foundation, Paavo Koistinen Foundation (AL.L. lab), Sigrid Juselius Foundation and Aatos Erkko Foundation (Grant number 210013) (T.K. lab). The study benefited from samples from Auria Biopankki Turku, Finland.

## Author contributions

Conceptualization, AL.L., P.P. and J.H.; Supervision, AL.L. and M.H.; methodology, P.P., J.H. and EY.D.; Software, P.P. and J.H.; Validation, T.K., M.A., EM.T, P.T., M.K. and T.E.; Formal analysis, J.H. and P.P.; Investigation, J.H., P.P., A.J.D. I.S. and HR.T.; Resources, AL.L., M.H., M.U.K., M.K, M.H. and T.K.; Data curation, P.P. and J.H.; Writing - Original Draft, J.H., AL.L., and P.P.; Writing - Review & Editing, M.H., T.K., A.J.D. and M.U.K.; Visualization, J.H., P.P. and A.J.D.; Project administration, AL.L.; Funding acquisition, AL.L and M.H.

## Declaration of interests

The authors declare no competing interests.

## Supplemental information

**Figure S1.**
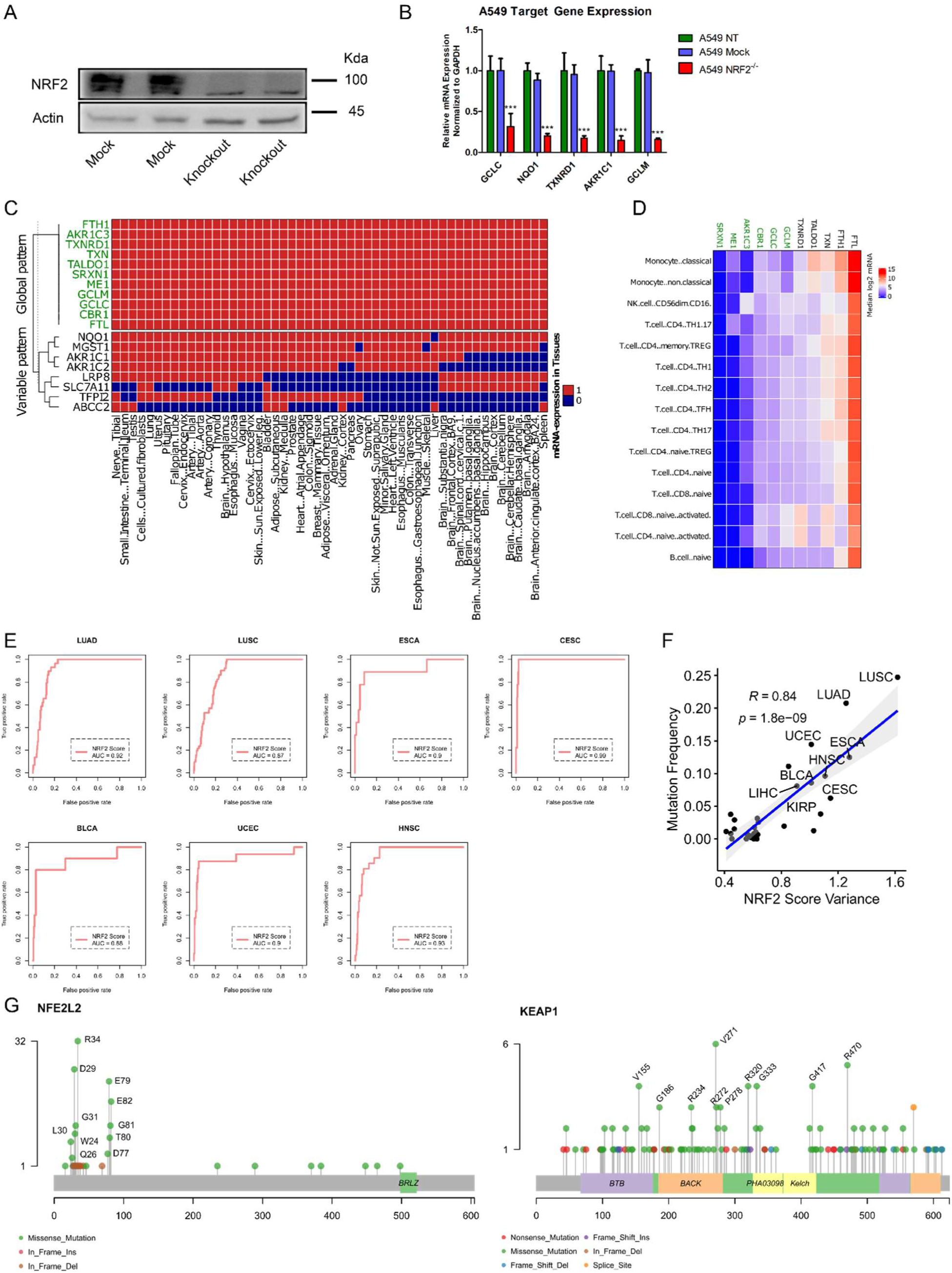
**A:** Western blot for A549-NRF2 knockout vs wild type. **B:** RT-qPCR for A549 knockout vs wild type (mock) and parental (NT). **C:** Binary expression patterns (median log2 CPM > 1.5) of score NRF2 target gene candidates. Genes passing the threshold across the dataset are colored green. **D:** Median gene expression in DICE database. Genes selected as the final score are colored green. **E:** Benchmark ROC of the NRF2 score against functional somatic *NFE2L2*/*KEAP1* mutations in cohorts with > 5 % *NFE2L2*/*KEAP1* mutation frequency. *SINGH NFE2L2 targets* was used as the reference gene set. **F:** Scatter plot of NRF2 score variance against *NFE2L2*/*KEAP1* mutation frequency. **G:** Lollipop plots for mutations in *NFE2L2* and *KEAP1* in TCGA NRF2 hyperactive cases classified by the NRF2 score (TPR > 0.85).

**Figure S3.**
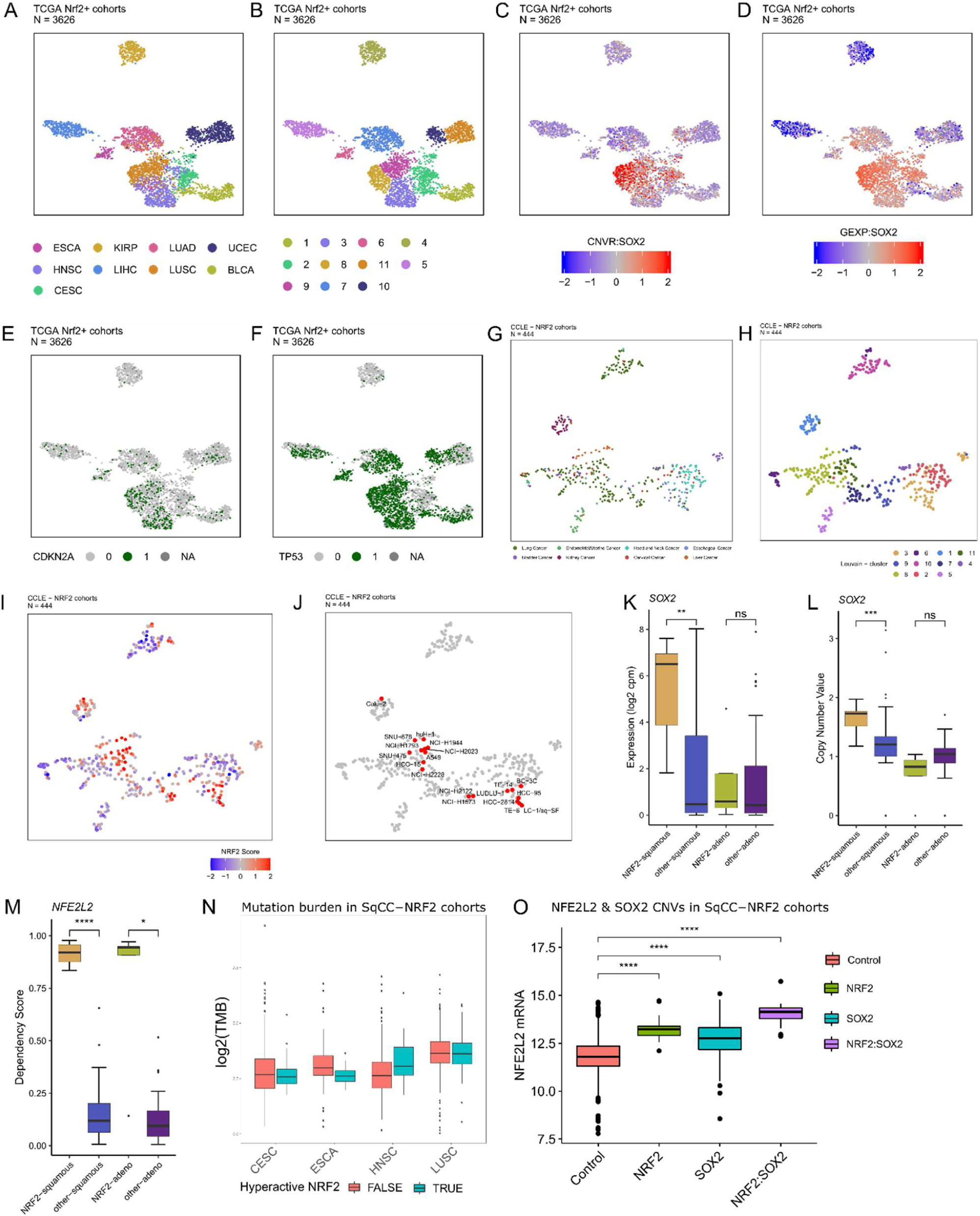
**A:** UMAP representation of TCGA NRF2 cohorts. **B:** UMAP representation of identified TCGA clusters. **C:** UMAP representation of *SOX2* copynumber. **D:** UMAP representation of *SOX2* gene expression. **E:** UMAP representation of *CDKN2A* mutation. **F:** UMAP representation of *TP53* mutation. **G:** UMAP representation of CCLE diseases. **H:** UMAP representation of identified CCLE clusters. **I:** UMAP representation of NRF2 score in CCLE. **J:** UMAP representation of CCLE cell-lines. **K:** Boxplot for SOX2 mRNA in CCLE data. **L:** Boxplot for SOX2 copy number in CCLE data. **M:** Boxplot for NFE2L2 dependency scores in CCLE data. The dependency score is interpreted as probability of dependency from 0 to 1. **N:** Mutation burden in TCGA SqCC cohorts stratified by NRF2 hyperactivity. **O:** *NFE2L2* and *SOX2* CNVs effect on *NFE2L2* mRNA in SqCC cohorts.

**Figure S4.**
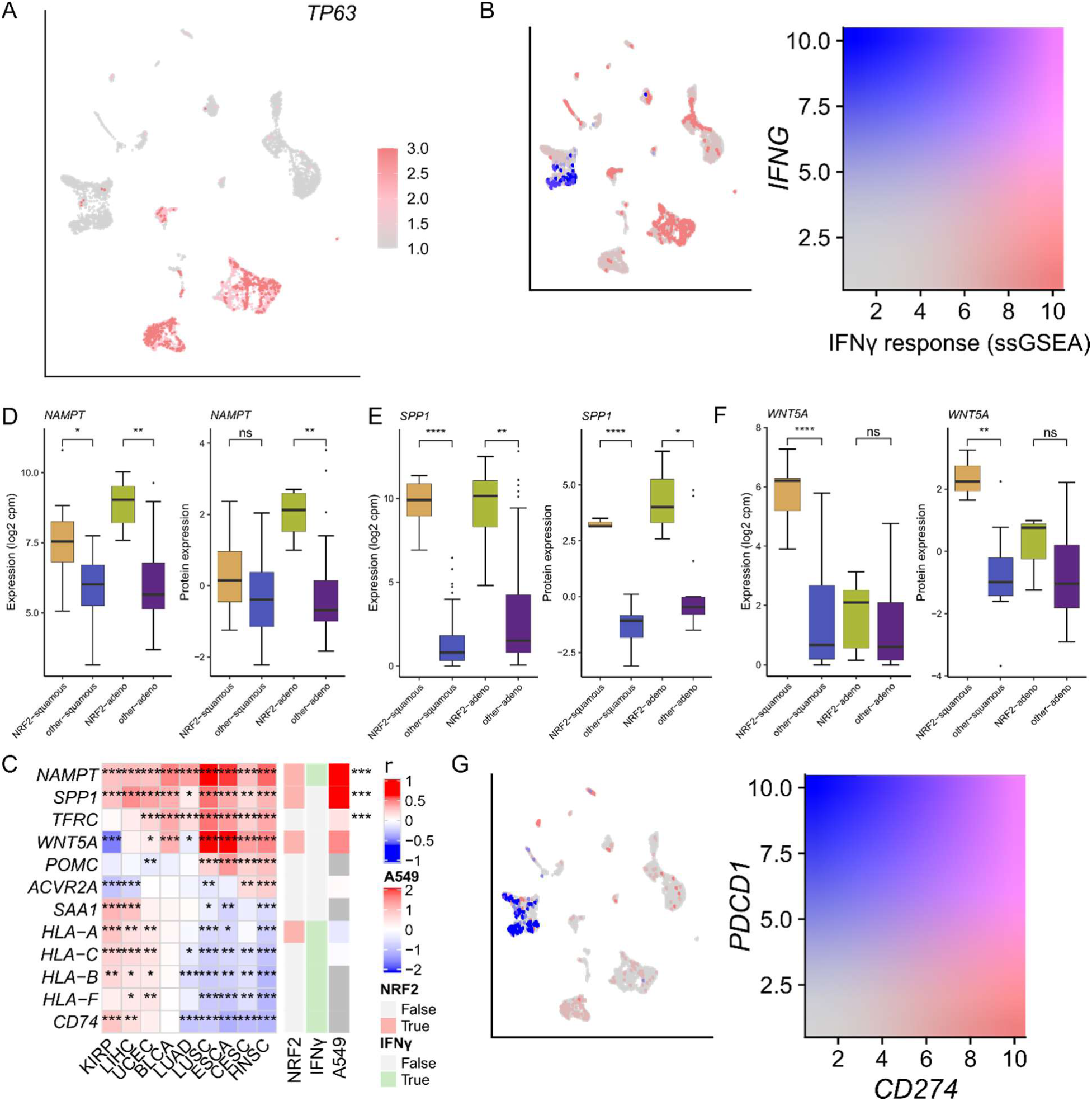
**A:** TP63 expression across the single cell populations. **B:** IFNγ response (ssGSEA) and *IFNG* expression across the single cell populations. **C:** TCGA gene expression to NRF2 score correlations for hits from the single-cell analysis. The NRF2 catalogue hits, interferon gamma associated genes and A549^OEvsKO^ differential expression are shown in the right annotation panel from left to right, respectively. **D:** NAMPT mRNA and protein expression in CCLE data in the identified adenocarcinoma and SqCC NRF2 clusters. **E:** SPP1 mRNA and protein expression in CCLE data in the identified adenocarcinoma and SqCC NRF2 clusters. **F:** WNT5A mRNA and protein expression in CCLE data in the identified adenocarcinoma and SqCC NRF2 clusters. **G:** PD-L1 (*CD274*) and PD-1 (*PDCD1*) expression across the single cell populations.

**Figure S5.**
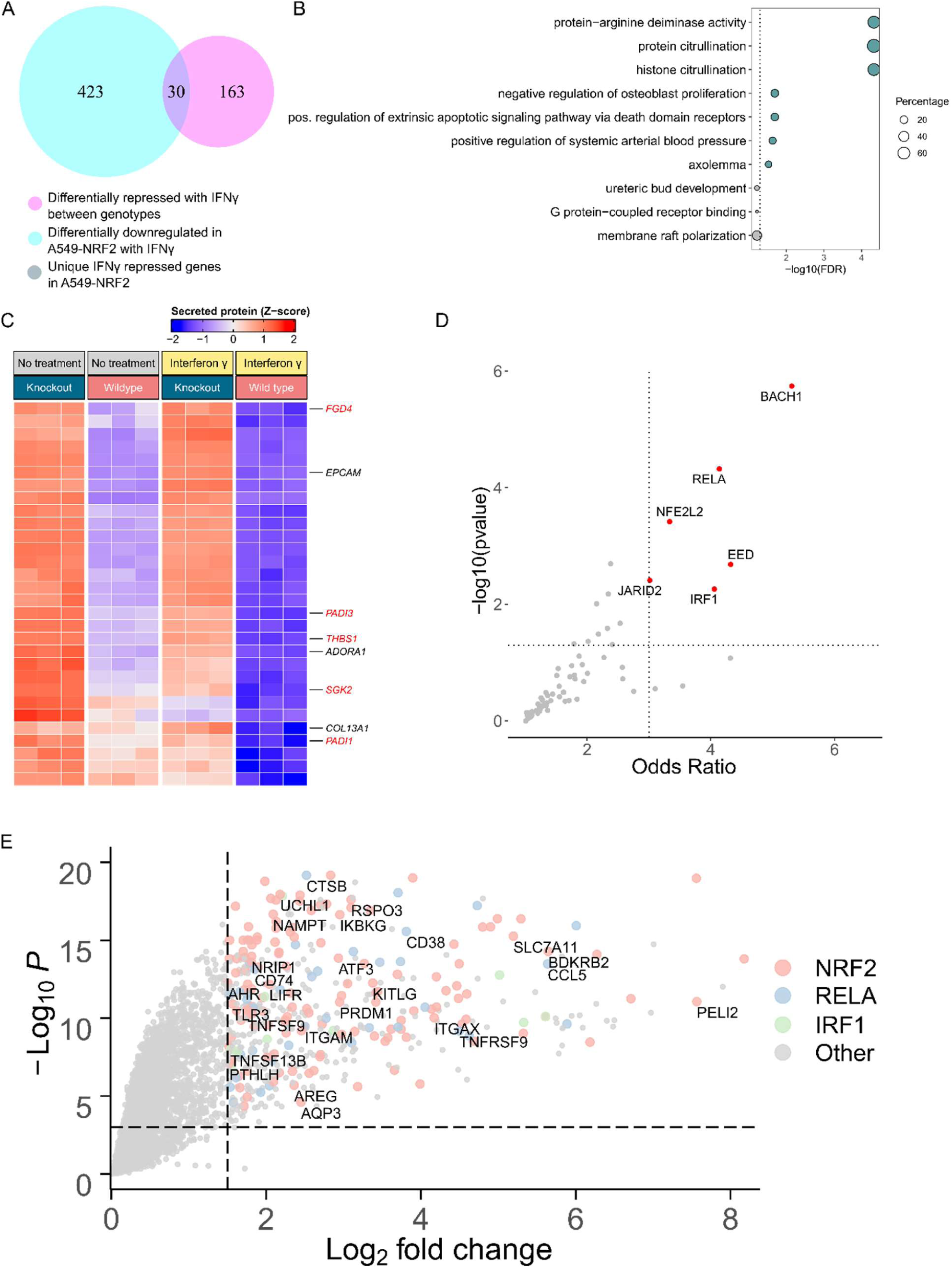
**A:** Venn-diagram of genes differentially downregulated in A549-NRF2^OEvsKO^ and differentially repressed between genotypes with IFNγ. **B:** Gene set enrichment analysis (GeneSCF) for the IFNγ repressed genes downregulated in A549-NRF2^OEvsKO^ with IFNγ treatment. **C:** Heatmap of IFNγ repressed genes downregulated in A549-NRF2^OEvsKO^ with IFNγ treatment. Immunogenomic terms and putative NRF2 targets (colored red) are annotated on the right. **D:** Transcription factor overrepresentation in unique IFNγ induced genes (overlapping genes in Figure 5a) of NRF2 hyperactive A549. **E:** Differentially upregulated genes downstream of NRF2, RELA & IRF1 in A549-NRF2 with IFNγ (immunomodulatory genes from TISIDB & InnateDB are labeled).

**Figure S6.**
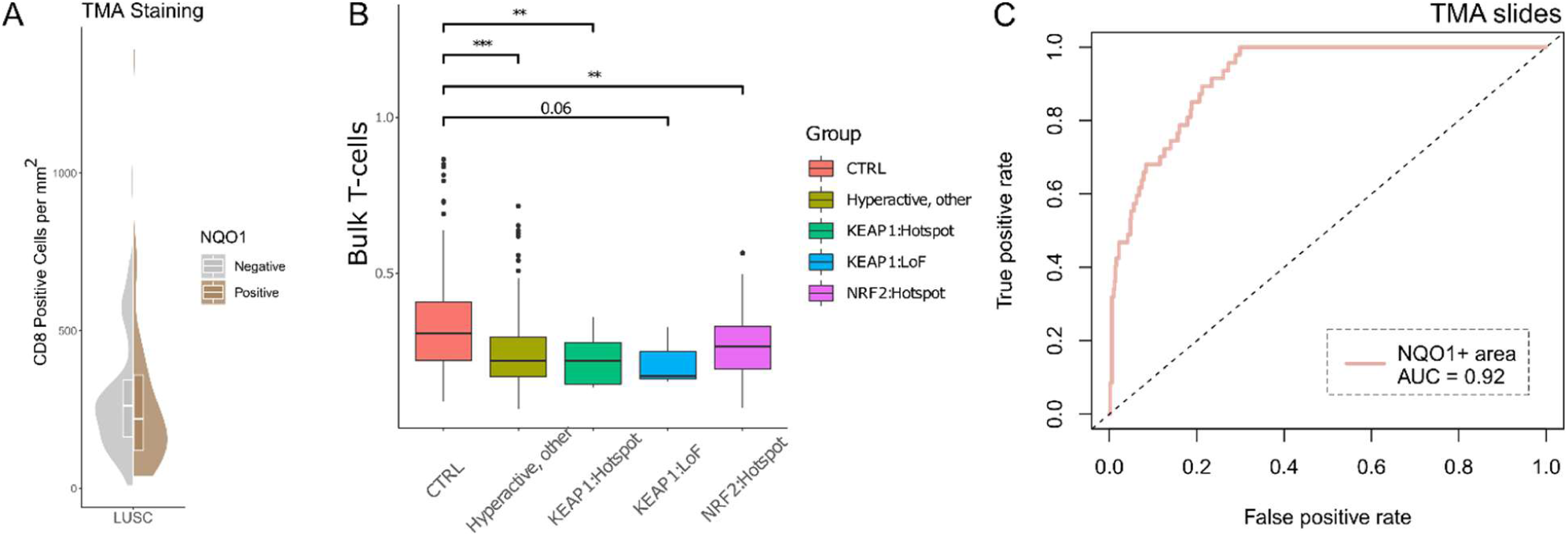
**A:** Relationship of CD8 positive cells and NQO1 staining (P = 0.09) in LUSC. **B:** *NFE2L2* and *KEAP1* mutations and bulk T-cell content in TCGA LUSC cohort. **C:** Benchmark ROC of the sample total NQO1 area to human assessed NQO1 status.

### Supplementary tables

Table 1: A549-NRF2^OEVSKO^ RNAseq (.xlsx)

Table 2: NRF2 candidate target genes (.xlsx)

Table 3: NRF2 score values in TCGA samples (.xlsx)

Table 4: Integrated GSEA (.xlsx)

Table 5: SOX2 candidate target genes (.xlsx)

Table 6: TP63 candidate target genes (.xlsx)

## Materials and Methods

### Processing of genome-wide multilevel data

#### TCGA

Processed multiomics data, sample level analysis results and clinical data, were retrieved from https://gdc.cancer.gov/about-data/publications/panimmune (Thorsson et al 2018) and https://gdc.cancer.gov/node/905/ (PanCanAtlas) for 33 cancer types available in TCGA cohort.

#### CCLE

Data was downloaded from https://portals.broadinstitute.org/ccle/data and https://depmap.org/portal/.

### Functional genomics analysis of NRF2 target genes

#### Enhancer and promoter catalogue generation

A549 and ENCODE genomic data for other cell types were used as reference to generate broad catalogue of candidate regulatory elements, that could be linked with NRF2 regulation. DNAse hotspots and ChromHMM classified genomic sites were downloaded from ENCODE for A549. Furthermore, DNAse clustered sites version 3 were downloaded from ENCODE for all cell types. Promoters were defined primarily based on A549 ChromHMM annotation, but if the annotation was missing from the data, promoters were defined as 500 bp upstream and 1500 bp downstream from each transcription start sites. Next, A549 DNAse hotspot sites were integrated with ENCODE DNAse clusters for various cell types, with score above 400 and cluster identified in at least 20 cell types. Promoters were intersected with DNAse hotspots to identify regulatory sites at promoters.

#### GRO-seq regression analysis of enhancer elements

eRNA can be used to detect active enhancers and it correlates with the gene expression of the target gene (Bouvy-Liivrand *et al*., 2017). To infer enhancer-gene pairs, we utilized publicly available GRO-seq data curated from GEO (GSE51225, GSE51633, GSE53964, GSE60454, GSE62046, GSE62296, GSE52642, GSE66448, GSE84432, GSE67519, GSE67540, GSE101803, GSE96859, GSE102819, GSE86165, GSE91011, GSE67295, GSE154427, GSE136813, GSE118530, GSE94872, GSE92375 and GSE117086). The final dataset comprised 336 samples representing 45 cell types. Homer analyzeRepeats.pl software was used for GRO-seq quantification of nascent RNA gene expression using gene introns as coordinates, with parameters: -strand + -noadj -noCondensing -pc 3. For enhancers, in case of intragenic enhancers, quantification was performed from the opposite strand and for intergenic enhancers from both strands, using the same parameters as before. Enhancer start and end coordinates were expanded by 500bp. Gene end coordinates were expanded by 5000bp for annotating intragenic gene enhancers and account for transcription at the end of transcripts. A linear model was fitted for each gene-enhancer pair in the same TAD that had Spearman’s Rho > 0 and P-value < 0.05. R^2^ value was used to estimate, how much of the gene expression could be explained by the eRNA expression.

#### NRF2 target gene identification

To infer transcription factor binding at the enhancer sites, ChIP-seq data for NFE2L2. The enhancer-gene pairs were defined as features with regression P-value < 0.05 and R^2^ > 0.1, and the maximum distance was set to 500kbp. To infer NRF2 bound regions, the preprocessed ENCODE ChIP-seq data for *NFE2L2* was downloaded from GEO (Datasets GSE91997, GSE91894, GSE91809, GSE91565). The ARE-motif from JASPAR database (*NFE2L2*, MA0150.1) was queried across the ChIP-peaks with Homer annotatePeaks, and NRF2 motif score cutoff for a binding site was set to 6. From these data, the minimum amount of ChIP-seq datasets with the given peak was set to two. Using this analysis framework 3000 bidirectional and 2000 intragenic enhancers were identified for NRF2.

### Development of the NRF2 score

A computational NRF2 score was developed by utilizing multiple datasets. The final gene set was derived by applying the following rationale: a) ChIP-seq peaks must be present at ChromHMM-promoters across the ENCODE *NFE2L2* ChIP-seq data b) the gene must be differentially expressed in A549-NRF2^OEvsKO^ (log2 fold-change > 1.5, Bonferroni adjusted p < 1e-5); and c) mRNA-expression must be observed across tissues in GETxportal data (https://www.gtexportal.org/home/datasets) and expression must be uniform across cell types in DICE-database (https://dice-database.org/), to counter variance arising from tissue specific expression and the immune-milieu, respectively.

Differential expression analysis for A549-NRF2^OEVSKO^ was conducted with limma. Direct NRF2 targets were defined as having a NRF2 binding in the NRF2 ChIP-seq data in the ChromHMM promoters: the peaks were narrowed down to regions that intersect all four datasets and include an ARE-sequence (motif score > 6).

Genes with tissue-specific expression patterns were excluded based on the expression profiles in the GETxportal data: median tissue expression cutoff was set to 1.5 log2 TPM for binary filtering (Figure S1A). Genes exhibiting high values or observable variance in median expression in immune-cells were excluded heuristically (Figure S1B).

The genes passing all of the aforementioned steps were defined as the NRF2 signature. The samples were scored with a geometric mean from the genes linear mRNA-expression. For TCGA samples, the score was normalized within each disease to the peak of the score distribution (mode). Performance of the scores was evaluated with receiver operating characteristic against functional NRF2 activating somatic variants (defined in OncoKB) using the ROCR-package for R.

### Gene Set Enrichment Analysis

GSEA (Java-GSEA) was performed for the selected TCGA-cohorts and CCLE cell lines of the matching diseases using NRF2 score as a continuous feature. Pan-cancer analysis for TCGA cancer types was conducted by calculating Stouffer’s statistic for each gene set across the selected cohorts. For the A549 cells, GSEA was conducted with a binary feature of NRF2^-/-^ vs NRF2^+/+^. Multiple-comparison corrections were performed with the Benjamini-Hochberg (BH) –method.

### UMAP analysis and Louvain clustering of TCGA and CCLE data

UMAP dimensionality reduction analysis was performed for 750 most variable genes using uwot R package, with parameters set to n.neighbors=12 and min.dist=0.4. Community detection based clustering for the same 750 most variable genes was performed using the Louvain algorithm, with k=4 implemented in igraph R package. Similarly, for CCLE 750 most variable genes were used with n.neighbors=6 and min.dist=0.3 and k=4. Various number of most variable genes were used for UMAP to confirm robustness of sample group detection.

### Functional genomics analysis of SOX2 and TP63 target genes

Target genes for SOX2 and TP63 were inferred from ChIP-seq data of the respective transcription factors (GEO dataset GSE46837). All parameters were set as in the analysis for NRF2 target genes (*NRF2 target gene identification*) using the enhancer and promoter catalogues defined earlier.

### Differential expression analysis for the NRF2 squamous cluster

Fold-differences across the transcriptome were computed between the NRF2 squamous cluster vs other squamous samples, and the statistical significance of differentials of gene-expression distributions were assessed with two-sided Wilcoxon tests, and subsequently multiple-comparison corrected with the BH-method. The result was filtered with FDR < 0.001 and a logFC threshold of 1 for absolute values. The overlap of the resulting genes were assessed with a) CCLE gene-expression and protein correlation to NRF2-score; b) genes in CellphoneDB and TISIDB (literature annotation and CRISPR-screen resistance genes); and c) the candidate target gene catalogues of NRF2, TP63 and SOX2. High-confidence hits (shown in Figure 3E) were defined as genes that correlate in CCLE protein or mRNA (FDR < 0.1, R > 0.25), are listed in CellPhoneDB or TISIDB and are listed in the target gene catalogues for at least one of the transcription factors.

### *In vitro* experiments

#### Cell culture

Cells were incubated in 37°C and 5 % CO_2_ throughout the experiments and the passage number was kept under 10 over the course of this study. Cell passaging was conducted before reaching confluency.

#### CRISPR/Cas9 NRF2 knockout

A549 cells were transfected with a 20bp NRF2 targeting (exon 4 of NFE2L2 ENST00000397062.7, CAAGCTGGTTGAGACTACCA) single-guide-RNA and scramble-sequence including plasmid vectors (SpCas9(BB)-2A-GFP (Addgene, PX458). Nrf2 knockout and control cells were generated via CRISPR-Cas9-mediated non-homologous end-joining (NHEJ). Transfection-positive cells were sorted with fluorescence-activated cell sorting (FACS) to obtain clonal populations. Validation of the clones was conducted with NRF2 western blot (Figure S1A), Sanger sequencing (Data not shown) and RT-qPCR (Figure S1B).

#### IFNγ treatment

A549 cells were seeded in 6-well plates and serum starved for 6 h with 0.5% FBS containing growth media. Cells were then treated with 10 ng/ml of IFNγ (catalog no. 300-02, Peprotech, USA) for 24 h and RNA extraction from the treated cells was done using High Pure RNA Isolation Kit (Roche, Lot#: 47931400). RNA samples were dissolved in nuclease-free water and stored in - 70° C until use.

#### Sanger sequencing and western blot

DNA was extracted with Genejet Genomic DNA purification kit (Thermo Scientific, catalog. no: K0702) and using the GENEWIZ Sanger Sequencing Services.

Cells were lysed and protein concentration measured using BCA kit (Pierce). 30 µg of total protein with 1X SDS-PAGE sample buffer (Biorad) was loaded in 4-20% mini-Protean TGX gels (Biorad) and the gel electrophoresis was done using Tris glycine running buffer containing SDS. Proteins were then transferred onto nitrocellulose membrane (0.2 µm, Biorad) using Owl Hep-1 semi-dry transfer system (Thermo scientific) following manufacturer’s instructions. The blots were blocked with 5% milk-TBST solution for 1 h at room temperature (RT). Blots were stained overnight at 4° C with NRF2 (1:5000 dilution, catalog no.16396-1-AP, Proteintech) and beta-actin (1:5000 dilution, catalog no. sc-47778, Santa Cruz, USA) antibodies in 5% milk-TBST solution. And secondary staining was done for 1 h at RT using Alexafluor 488-or 680-labelled anti-rabbit and anti-mouse antibodies (Invitrogen) in 2% milk-TBST solution. The blots were then visualized using Biorad documentation system.

#### RNAseq & differential expression analysis

RNAseq was conducted at FIMM genomics NGS transcriptomics service (University of Helsinki) aiming at 30 million reads per sample with 100 bp read length. Alignment was performed with Hisat2 and differential expression analysis was conducted with the limma R-package using the standard protocol from the user manual. (https://www.bioconductor.org/packages/devel/bioc/vignettes/limma/inst/doc/usersguide.pdf) For IFNγ treatment, two separate differential expression analyses were conducted: a) differential expression between one factor - genotype - over the interferon-gamma treatment; and b) a model with two factors - genotype and treatment - was used to assess the genes differentially induced by IFNγ between genotypes (ΔNRF2^+/+^_IFNγ-DMSO_ – ΔNRF2^-/-^_IFNγ-DMSO_). The differentially expressed genes affected by interferon-gamma were defined as genes passing the following criteria: p < 0.01 & logFC > 1.5 in analysis a and p < 0.01 & logFC > 0.5 in analysis b, that is, induction would *de facto* result in pronounced differential absolute expression with IFNγ.

#### Proteome profiler

Cells were seeded at 150000 cells per well in 12 well plates, incubated overnight and serum deprived the following day ± 10 ng/ml IFNγ (catalog no. 300-02, Peprotech, USA). Upon 48 h of IFNγ administration, the media was collected, centrifuged at 500g for 5 min, and processed according the Proteome Profiler XL Cytokine Array Kit (R&D Biosystems, catalog no. ARY022B) instructions with overnight primary antibody incubation at +4°C. The blots were imaged with Bio-Rad ChemiDoc MP Imaging System. The data was analyzed with ImageJ v. 1.47, normalizing mean intensity values of the dots to internal background and positive control. Students T-test was used to statistically assess group differences.

### Copy-number analysis

Copy-number analysis was conducted for firehose-derived TCGA segment-files for HNSC, LUSC, ESCA, and CESC, stratified by the NRF2 subtype (cluster 8) with Gistic2 using the following parameters: *-ta 0.1 -Peakpeel 1 -brlen 0.7 -cap 1.5 -conf 0.99 -td 0.1 -genegistic 1 –gcm extreme -js 4 -maxseg 2000 -qvt 0.25 -rx 0 -savegene 1 -broad 1*.

### Processing of single cell data

The HNSC single cell dataset was downloaded from GEO (GSE103322) and processed with Seurat R-package v3. Cells with more than 8,000 detected genes were filtered out. Seurat SCTransform with 3,000 variable features was used for data normalization. 25 principal components and default parameters were used for UMAP projection and Louvain clustering. SingleR 1.0.1 (Aran *et al*., 2019) was used for the automated cell type annotation. These annotations were manually refined according to cell types identified in Puram et al., 2017.

### CellPhoneDB analysis

The HNSC single-cell dataset was analyzed with CellPhoneDB statistical analysis method using 1000 iterations and an expression threshold of 0.1. To assess the differential interactions, the data was visualized as a ratio of interactions means in the NRF2 cluster vs other cancer clusters, only including interactions that were exclusively statistically significant in either the NRF2 cluster (positive ratio) or the other clusters (negative ratio). Statistically non-significant ratios were removed from the heatmap for clarity.

### Survival analysis

A pan-HNSC-NSCLC-SKCM dataset was downloaded from GEO (GSE110390) and the associated clinical data was obtained from TIDE-database (http://tide.dfci.harvard.edu/). SKCM samples were excluded from the analysis, and the nCounter PanCancer Immune Profiling Panel targets were normalized to 40 housekeeping genes with NanoStringNorm R-package. Mean normalized log2 mRNA expression was used as a cutoff to dissect *SPP1* high and low expressing samples. Survival analysis was performed with survival R-package. The Kaplan-Meier curve was plotted with survminer R-package.

### CIBERSORT analysis for TCGA data

CIBERSORT was run for the TCGA data with the CIBERSORT R-package separately for each cancer type with the absolute method (*sig.score*), using 100 permutations and without quantile normalization. Correlations to NRF2 score were assessed with Pearson’s method. The bulk lymphocyte analysis was conducted by taking the sum of all T-lymphocyte fractions, and using a cutoff for NRF2 hyperactivity as TPR > 0.85 from the ROC-analysis. The statistics were computed with Mann-Whitney U.

### Immunohistochemistry

NSCLC tissue microarray sections (LUAD n = 211, LUSC n = 117) were obtained from Auria Biobank, Turku University Hospital, Turku, Finland. NQO1 (Cell Signaling NQO1 A180 Mouse mAb, 1:250; catalog. no. 3187S) was stained by incubating overnight at +4C°. NQO1 signal was quantified as the mean of relative positive-stained area across replicates, and NQO1-positive samples were defined as Q_3_ of the mean NQO1-signal across biological replicates. The NQO1 area quantification was benchmarked to a human evaluated assessment of binary NQO1 positivity of a sample (Figure S6B). For T lymphocyte immunohistochemistry, TMA-sections were stained with anti-CD3 (LN10, 1:200; Novocastra) and anti-CD8 (SP16, 1:400; Thermo Scientific) using a LabVision Autostainer 480 (ImmunoVision Technologies Inc.). Antigen retrieval was done with Tris-EDTA buffer at pH 9 by microwaving the slides in 98 degrees Celsius for 15 minutes. Samples were incubated with diluted antibodies for 30 minutes at room temperature. Diaminobenzidine (DAB) was used as a chromogen and haematoxylin as a counterstain. Positive control tissue for CD3 and CD8 immunohistochemistry was normal tonsil. The slides were digitized with a slide scanner (Nano Zoomer XR, Hamamatsu) and quantification of CD3+ and CD8+ T cells was performed using QuPath, an open-source bioimage analysis software (version 0.1.2) (Bankhead, P., Loughrey, M.B., Fernández, J.A. *et al*. QuPath: Open source software for digital pathology image analysis (Bankhead *et al*., 2017). With the a priori assumption of a negative relationship between the variables based on earlier observations, one-sided Mann-Whitney U test was used for statistical inference between groups.

## Notes

### Competing Interest Statement

The authors have declared no competing interest.

## References

Ancevski Hunter, K., Socinski, M.A. and Villaruz, L.C. (2018) “PD-L1 Testing in Guiding Patient Selection for PD-1/PD-L1 Inhibitor Therapy in Lung Cancer,” Molecular Diagnosis and Therapy, 22(1). doi:10.1007/s40291-017-0308-6.

Aran, D. et al. (2019) “Reference-based analysis of lung single-cell sequencing reveals a transitional profibrotic macrophage,” Nature Immunology, 20(2). doi:10.1038/s41590-018-0276-y.

Bankhead, P. et al. (2017) “QuPath: Open source software for digital pathology image analysis,” Scientific Reports, 7(1). doi:10.1038/s41598-017-17204-5.

Best, S.A. et al. (2018) “Synergy between the KEAP1/NRF2 and PI3K Pathways Drives Non-Small-Cell Lung Cancer with an Altered Immune Microenvironment,” Cell Metabolism, 27(4). doi:10.1016/j.cmet.2018.02.006.

Bouvy-Liivrand, M. et al. (2017) “Analysis of primary microRNA loci from nascent transcriptomes reveals regulatory domains governed by chromatin architecture,” Nucleic Acids Research, 45(17). doi:10.1093/nar/gkx680.

Breuer, K. et al. (2013) “InnateDB: Systems biology of innate immunity and beyond - Recent updates and continuing curation,” Nucleic Acids Research, 41(D1). doi:10.1093/nar/gks1147.

Campbell, J.D. et al. (2016) “Distinct patterns of somatic genome alterations in lung adenocarcinomas and squamous cell carcinomas,” Nature Genetics, 48(6). doi:10.1038/ng.3564.

Chakravarty, D. et al. (2017) “OncoKB: A Precision Oncology Knowledge Base,” JCO Precision Oncology [Preprint], (1). doi:10.1200/po.17.00011.

Duan, Q. et al. (2020) “Turning Cold into Hot: Firing up the Tumor Microenvironment,” Trends in Cancer. doi:10.1016/j.trecan.2020.02.022.

Efremova, M. et al. (2020) “CellPhoneDB: inferring cell–cell communication from combined expression of multi-subunit ligand–receptor complexes,” Nature Protocols, 15(4). doi:10.1038/s41596-020-0292-x.

Granot, Z. (2019) “Neutrophils as a Therapeutic Target in Cancer,” Frontiers in immunology. doi:10.3389/fimmu.2019.01710.

Hanahan, D. and Weinberg, R.A. (2011) “Hallmarks of cancer: The next generation,” Cell. doi:10.1016/j.cell.2011.02.013.

Hayes, J.D. and Dinkova-Kostova, A.T. (2014) “The Nrf2 regulatory network provides an interface between redox and intermediary metabolism,” Trends in Biochemical Sciences. doi:10.1016/j.tibs.2014.02.002.

Hayes, J.D., Dinkova-Kostova, A.T. and Tew, K.D. (2020) “Oxidative Stress in Cancer,” Cancer Cell. doi:10.1016/j.ccell.2020.06.001.

Hsieh, M.H. et al. (2019) “p63 and SOX2 Dictate Glucose Reliance and Metabolic Vulnerabilities in Squamous Cell Carcinomas,” Cell Reports, 28(7). doi:10.1016/j.celrep.2019.07.027.

Ivashkiv, L.B. (2018) “IFNγ: signalling, epigenetics and roles in immunity, metabolism, disease and cancer immunotherapy,” Nature Reviews Immunology. doi:10.1038/s41577-018-0029-z.

Kansanen, E. et al. (2013) “The Keap1-Nrf2 pathway: Mechanisms of activation and dysregulation in cancer,” Redox Biology. doi:10.1016/j.redox.2012.10.001.

Klement, J.D. et al. (2018) “An osteopontin/CD44 immune checkpoint controls CD8+ T cell activation and tumor immune evasion,” Journal of Clinical Investigation, 128(12). doi:10.1172/JCI123360.

Lachmann, A. et al. (2010) “ChEA: Transcription factor regulation inferred from integrating genome-wide ChIP-X experiments,” Bioinformatics, 26(19). doi:10.1093/bioinformatics/btq466.

Leinonen, H.M. et al. (2014) “Role of the keap1-Nrf2 pathway in cancer,” in Advances in Cancer Research. doi:10.1016/B978-0-12-420117-0.00008-6.

Liu, P. et al. (2021) “Nrf2 overexpression increases risk of high tumor mutation burden in acute myeloid leukemia by inhibiting MSH2,” Cell Death and Disease, 12(1). doi:10.1038/s41419-020-03331-x.

Lv, H. et al. (2021) “NAD+ Metabolism Maintains Inducible PD-L1 Expression to Drive Tumor Immune Evasion,” Cell Metabolism, 33(1). doi:10.1016/j.cmet.2020.10.021.

Newman, A.M. et al. (2015) “Robust enumeration of cell subsets from tissue expression profiles,” Nature Methods, 12(5). doi:10.1038/nmeth.3337.

Ottensmeier, C.H. et al. (2016) “Upregulated glucose metabolism correlates inversely with CD8+ T-cell infiltration and survival in squamous cell carcinoma,” Cancer Research, 76(14). doi:10.1158/0008-5472.CAN-15-3121.

Pölönen, P. et al. (2019) “Nrf2 and SQSTM1/p62 jointly contribute to mesenchymal transition and invasion in glioblastoma,” Oncogene, 38(50). doi:10.1038/s41388-019-0956-6.

Porter, L. and McCaughan, F. (2020) “SOX2 and squamous cancers,” Seminars in Cancer Biology. doi:10.1016/j.semcancer.2020.05.007.

Puram, S. v. et al. (2017) “Single-Cell Transcriptomic Analysis of Primary and Metastatic Tumor Ecosystems in Head and Neck Cancer,” Cell, 171(7). doi:10.1016/j.cell.2017.10.044.

Rojo de la Vega, M., Chapman, E. and Zhang, D.D. (2018) “NRF2 and the Hallmarks of Cancer,” Cancer Cell. doi:10.1016/j.ccell.2018.03.022.

Ross, D. and Siegel, D. (2017) “Functions of NQO1 in cellular protection and CoQ10 metabolism and its potential role as a redox sensitive molecular switch,” Frontiers in Physiology. doi:10.3389/fphys.2017.00595.

Ru, B. et al. (2019) “TISIDB: An integrated repository portal for tumor-immune system interactions,” Bioinformatics, 35(20). doi:10.1093/bioinformatics/btz210.

Saltz, J. et al. (2018) “Spatial Organization and Molecular Correlation of Tumor-Infiltrating Lymphocytes Using Deep Learning on Pathology Images,” Cell Reports, 23(1). doi:10.1016/j.celrep.2018.03.086.

Singh, A. et al. (2021) “NRF2 Activation promotes aggressive lung cancer and associates with poor clinical outcomes,” Clinical Cancer Research, 27(3). doi:10.1158/1078-0432.CCR-20-1985.

Spranger, S. and Gajewski, T.F. (2018) “Impact of oncogenic pathways on evasion of antitumour immune responses,” Nature Reviews Cancer. doi:10.1038/nrc.2017.117.

Subhash, S. and Kanduri, C. (2016) “GeneSCF: A real-time based functional enrichment tool with support for multiple organisms,” BMC Bioinformatics, 17(1). doi:10.1186/s12859-016-1250-z.

Tan, Y.S. et al. (2018) “Mitigating SOX2-potentiated Immune Escape of Head and Neck Squamous Cell Carcinoma with a STING-inducing Nanosatellite Vaccine,” Clinical Cancer Research, 24(17). doi:10.1158/1078-0432.CCR-17-2807.

Thorsson, V. et al. (2018) “The Immune Landscape of Cancer,” Immunity, 48(4). doi:10.1016/j.immuni.2018.03.023.

Wang, W. et al. (2019) “CD8+ T cells regulate tumour ferroptosis during cancer immunotherapy,” Nature, 569(7755). doi:10.1038/s41586-019-1170-y.

Wu, L. et al. (2019) “Tumor-associated neutrophils in cancer: Going pro,” Cancers, 11(4). doi:10.3390/cancers11040564.

Zhao, F. et al. (2018) “Paracrine Wnt5a-β-Catenin Signaling Triggers a Metabolic Program that Drives Dendritic Cell Tolerization,” Immunity, 48(1). doi:10.1016/j.immuni.2017.12.004.

Zhou, T. et al. (2020) “IL-18BP is a secreted immune checkpoint and barrier to IL-18 immunotherapy,” Nature, 583(7817). doi:10.1038/s41586-020-2422-6.

Zhu, B. et al. (2018) “Targeting the upstream transcriptional activator of PD-L1 as an alternative strategy in melanoma therapy,” Oncogene, 37(36). doi:10.1038/s41388-018-0314-0.

